# Deep molecular, cellular and temporal phenotyping of developmental perturbations at whole organism scale

**DOI:** 10.1101/2022.08.04.502764

**Authors:** Lauren M. Saunders, Sanjay R. Srivatsan, Madeleine Duran, Michael W. Dorrity, Brent Ewing, Tor Linbo, Jay Shendure, David W. Raible, Cecilia B. Moens, David Kimelman, Cole Trapnell

## Abstract

The maturation of single cell transcriptomic technologies has facilitated the generation of comprehensive cellular atlases from whole embryos. A majority of this data, however, has been collected from wild type embryos without an appreciation for latent variation present in development. Here we present single cell transcriptomic data from 1812 individually resolved developing zebrafish embryos, encompassing 19 time points, 23 genetic perturbations, and totaling 3.2M cells. The high degree of replication in our study (8 or more embryos per condition) allows us to estimate the variance in cell type abundance organism-wide and to detect perturbation-dependent deviance in cell type composition relative to wild type embryos. Our approach is sensitive to rare cell types, resolving developmental trajectories and genetic dependencies in the cranial ganglia neurons, a cell population that comprises less than 1% of the embryo. Additionally, time-series profiling of individual mutants identified a group of *brachyury*-independent cells with strikingly similar transcriptomes to notochord sheath cells, leading to new hypotheses about the origins of the skull. We anticipate that standardized collection of high-resolution, organism-scale single cell data from large numbers of individual embryos will enable mapping the genetic dependencies of zebrafish cell types, while also addressing long-standing challenges in developmental genetics, including the cellular and transcriptional plasticity underlying phenotypic diversity across individuals.

## Introduction

Understanding how each gene in our genome contributes to our individual phenotypes during embryogenesis is a fundamental goal of developmental genetics. Genetic screens in multicellular animals have enabled the dissection of diverse developmental processes, illuminating the functions of thousands of genes^1–4^. Although advances in automation, imaging, and genetic tools have increased the sophistication of phenotyping and yielded new insights into vertebrate development, phenotyping remains a significant bottleneck in characterizing gene function. Single-cell RNA-sequencing (scRNA-seq) applied at whole-embryo scale offers a comprehensive means of simultaneously measuring molecular and cellular phenotypes ^5–10^. However, realizing this promise requires overcoming several challenges: sequencing a very large number of cells through developmental time, rapidly generating mutant embryos, and sampling many individuals to account for biological variability during embryogenesis. These challenges have, until now, limited analyses to few genetic perturbations in comparatively less complex animals or at early stages of development.

Recent technological advances have created an opportunity to overcome these challenges, spurring a new era of developmental genomics. Combinatorial cellular indexing, or “sci-seq”, profiles the transcriptomes of millions of nuclei in one experiment, enabling embryo-scale analyses^6, 11, 12^. Labeling techniques that “hash” cells or nuclei from distinct samples allow one to multiplex specimens or whole embryos together^13, 14^, facilitating the analysis of many individuals. Parallel advances in CRISPR-Cas9 mutagenesis now enable programmatic, highly efficient genome editing at the F0 stage^15, 16^, circumventing the generation time required to create mutant embryos.

Here we describe the application of these three technologies to zebrafish, a model organism that develops rapidly, exhibits extensive cell type diversity, and is made up of a relatively small number of cells^7, 8, 17^. The data presented constitute two major efforts: 1) the establishment of an annotated, individually-resolved reference atlas, comprising 1,167 individuals and 1.2 million cells over 19 timepoints, filling a major gap in existing zebrafish atlases; and 2) the collection of perturbation data from 23 genetic perturbations over multiple timepoints, totaling 645 individuals and 2 million cells. By collecting many replicate embryos (8 or more embryos per condition), we implement statistical tests to systematically assess the gains and loss of cell types consequent to perturbation throughout the developing zebrafish. By comparing our harmonized reference and perturbation datasets, we dissect the genetic dependencies of rare cell types such as the the sensory neurons of the cranial ganglia, which comprise less than 1% of the cells in the organism. Finally, we leverage time-resolved, differential cell type-abundance analysis to characterize a cryptic population of cranial cartilage, explicating new hypotheses regarding the evolutionary origins of the vertebrate skull. Together, our scalable approach is flexible, comprehensive, cost-effective, and more uniform than conventional phenotyping strategies. We anticipate that this new experimental and analytical workflow will enable rapid, high-resolution phenotyping of whole developing animals to better understand the genetic dependencies of cell types in a developing organism.

### Time and individual-resolved cellular atlas of wildtype zebrafish development

To robustly detect perturbation-dependent changes in cellular composition, we adapted sci-Plex^13^, a workflow for multiplexing thousands of samples during single-cell RNA-seq, to barcode individual embryos and to capture single nucleus transcriptomes from whole organisms (**Methods**). We optimized whole embryo dissociations followed by oligo hashing to label each nucleus with an embryo-specific barcode, finding that we can unambiguously recover the embryo of origin for ∼70% of cells passing quality control thresholds (**Extended Data Fig. 1, Supplementary Table 1**).

**Figure 1.**
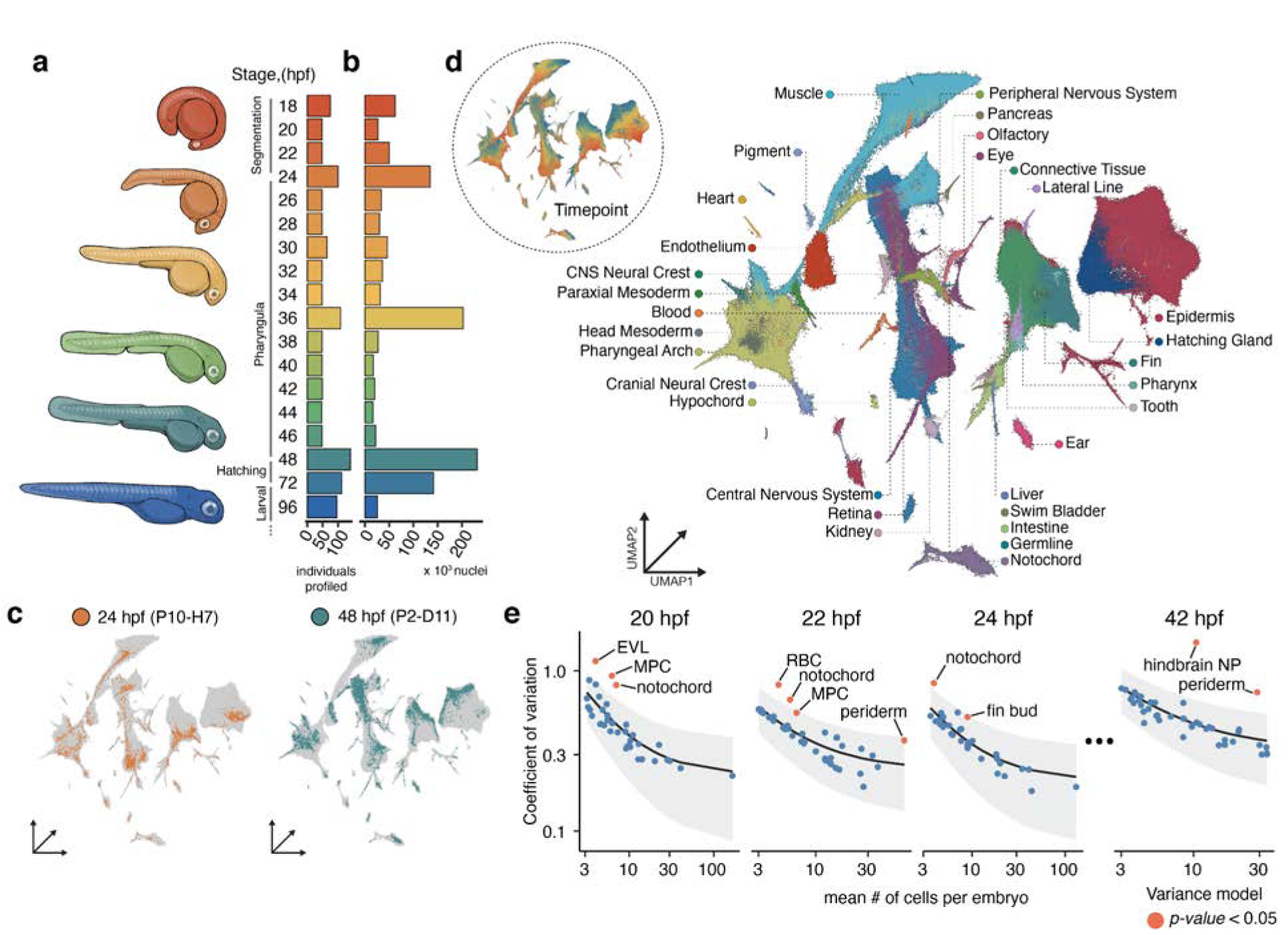
Collection of an individual-resolved single cell zebrafish atlas using oligo hashing. **a-b**, Number of individuals (**a**) and cells (**b**) profiled from each developmental time point. Representative drawings for select stages are shown (left) with colors matching timepoints in the bar graph. **c**, Cells originating from two individual embryos from 24 hpf (left) and 48 hpf (right) titled with the hash oligo barcodes. **d**, UMAP embedded in 3-dimensions, colored by tissue annotation. Inset colored by developmental time matching colors in panels a and b. **e**, Cell type count mean (x-axis) versus variance (y-axis) for a subset of timepoints. The coefficient of variation (black line) and standard error (gray fill) for each cell type’s abundance is modeled using a generalized linear model with a gamma-distributed response. Cell types that vary significantly more than expected relative to the model are colored in red (*p*-value < 0.05). (EVL - Enveloping layer; MPC - mesodermal progenitor cell; RBC - red blood cell; hindbrain NP - hindbrain neural progenitor (R7/8)).

Existing single cell atlases of zebrafish development document the emergence of diverse cell types from 3.3 hours (pre-gastrulation) to 5 days (late organogenesis) post fertilization, in addition to a few selected mutants at a single time point^7, 8, 17^ (**Extended Data Fig. 2**). While these datasets resolved diverse cellular states during zebrafish embryogenesis, each time point was a pool of embryos, thus masking heterogeneity between individuals. To assess variation resulting from gene knockouts, estimating the baseline heterogeneity present between individual wild type embryos is critical.

**Figure 2.**
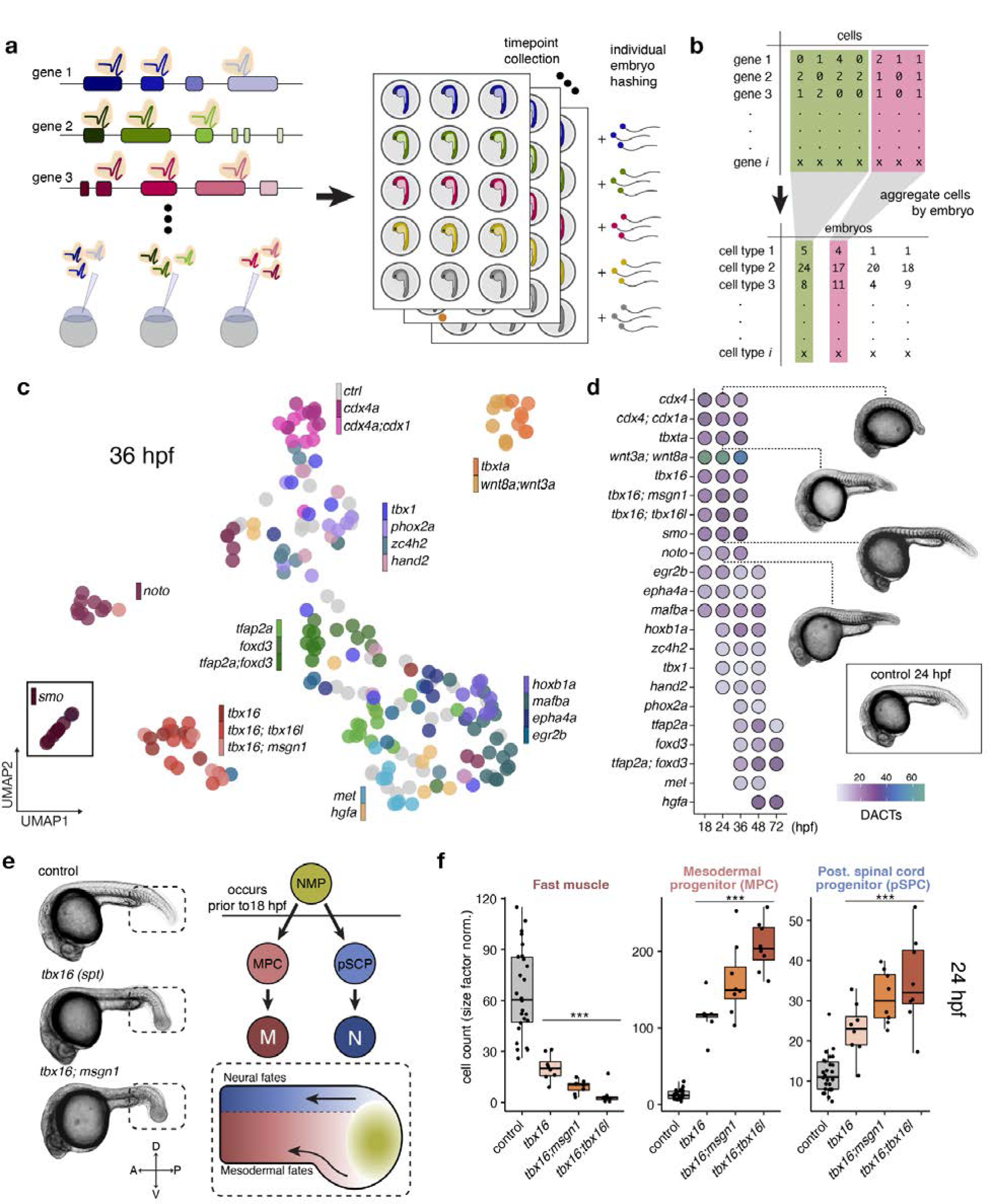
High resolution phenotyping of crispant (F0 CRISPR mutant) zebrafish embryos. **a**, A schematic of the experimental design. We designed 2-3 gRNAs across multiple exons and injected guide/Cas9 complexes at the one cell stage. Embryos were screened for desired phenotypes, dechorionated and dissociated in each well of a 96 well plate prior to nuclei isolation, hashing, and fixation. **b**, An individual by cell-type matrix was constructed by tallying the number of each broad cell type recovered for each individual embryo. **c**, UMAP embedding of individual composition data (as shown in panel b) at 36 hpf, embryos are colored by their genotype. *Smoothened* (*smo*) is shown as an inset because it was distant to the other embryos in the embedding. **d**, Heatmap of the number of differentially abundant cell types (DACTs) for each crispant, time point combination experiment. Cell types refers to the broad cell type annotation level (N = 85 total) and abundance differences were deemed significantly different if *q* < 0.01 (Beta binomial regression). Images are representative siblings of collected embryos at 24-26 hpf. **e**, Representative images of control, *tbx16* and *tbx16*;*msgn1* crispants at 24 hpf, accompanied by a schematic of mesodermal differentiation in the tail bud (dashed box). Neuromuscular progenitors (NMPs) give rise to two anteriorly migrating lineages of cells: 1) mesodermal progenitor cells (MPCs) and 2) posterior spinal precursor cells, which give rise to somitic muscle and spinal cord neurons, respectively. Compass rose denotes anatomical orientation: D-Dorsal, V-Ventral, A-Anterior and P-Posterior. **f**, Boxplots of cell counts from individual embryos across four cell types and four genotypes at 24 hpf. Significance (*** *q* < 1e-4) relative to control-injected embryos.

Moreover after late segmentation (18 hpf), intervals between sampling time points in these datasets were very sparse and therefore were not well resolved for key differentiation events during organogenesis. Thus, we first set out to establish a more high-resolution reference atlas with individual embryo resolution and fine-grained time point sampling.

We collected and labeled individual zebrafish embryos over 18 timepoints during embryonic and early larval development, spanning from 18 hours post fertilization (hpf), during late somitogenesis, with 2-hour resolution until 48 hpf, then a 72 hpf time point and 96 hpf time point, a period marking the early larval stages (**Fig. 1a**). After quality control, our dataset included ∼1.25 million cells from 1,223 barcoded individual embryos across 18 timepoints. At each timepoint, we collected between 48 and 140 embryos and amassed ∼17,000 to ∼231,000 high-quality, single nucleus transcriptomes per time point across four sci-RNA-seq3 experiments (**Fig. 1b,c, Extended Data Fig. 3**). These data also integrated coherently with published zebrafish scRNA-seq data from earlier and overlapping timepoints, despite collection on different platforms (**Extended Data Fig. 2b,c**). Cell type identity was inferred by inspection of marker genes for each cluster, which were cross-referenced with annotated gene expression data from the zebrafish genome database, ZFIN^18^. Overall, we hierarchically classified cells into 33 major tissues, 85 broad cell types, and 156 cell subtypes (**Fig. 1d, Extended Data Fig. 4, Supplementary Table 2**).

**Figure 3.**
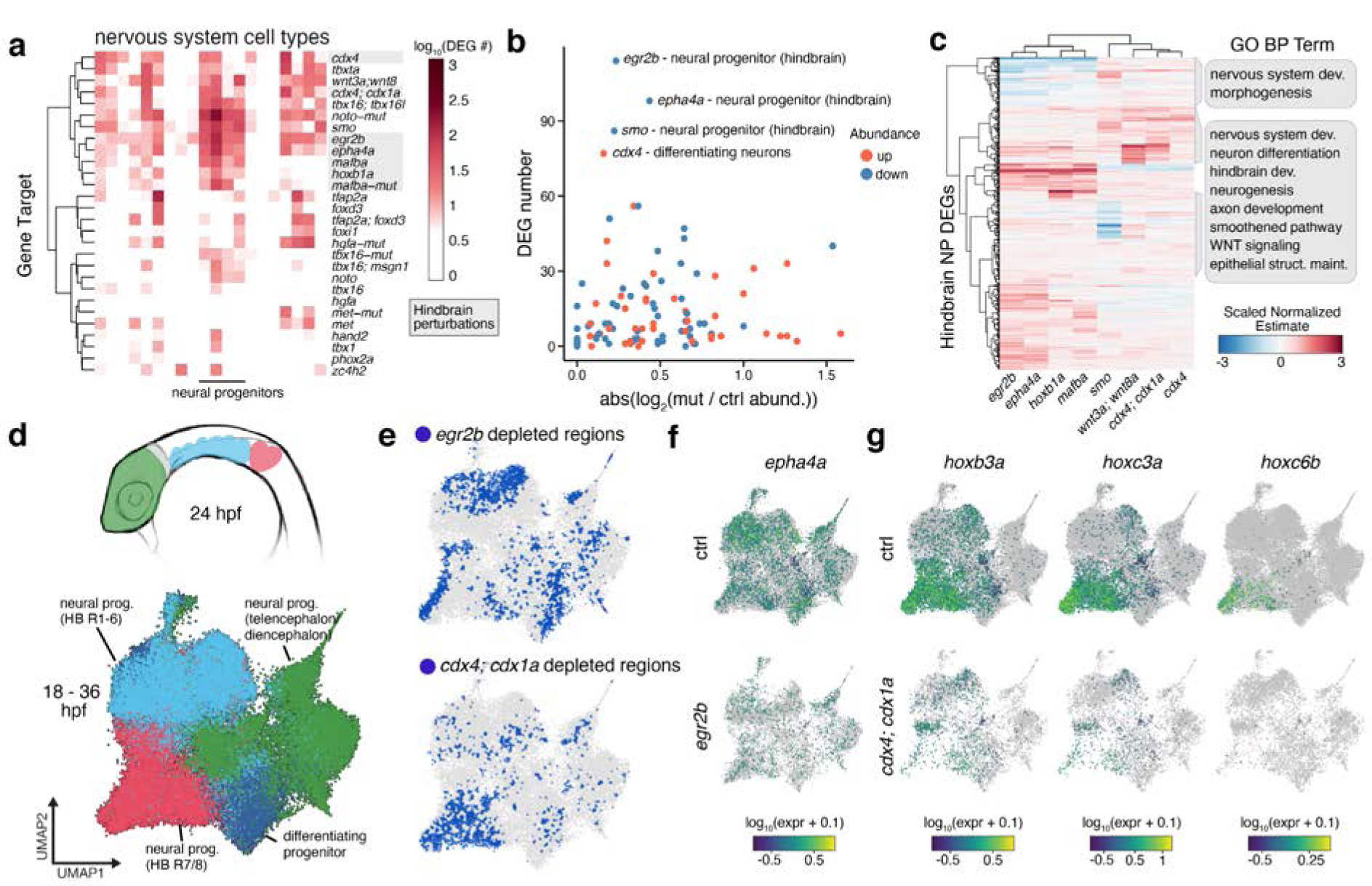
Systematic detection of differentially expressed genes (DEGs) and cell state changes across perturbations. **a**, Clustered heatmap displaying the number of DEGs (displayed as log_10_(x +1); *q*-value < 0.05) for neural cell typesxall perturbation combinations. Hindbrain perturbations are highlighted in gray. **b**, The number of DEGs vs. the absolute abundance change for hindbrain perturbation (highlighted in gray)xneural cell type combinations. All collected time points are shown with abundance change direction denoted by color. **c**, A heatmap of the DEG coefficient estimates for hindbrain neural progenitor cells of embryos from eight perturbations affecting hindbrain development. Select, significantly enriched Gene Ontology (Biological Process) termes are listed. **d**, (Top) Diagram of a 24 hpf zebrafish (anterior, lateral view), where anatomical regions are colored to match the UMAP embedding (bottom) of sub-clustered neural progenitors from all perturbations and timepoints. **e**, UMAP embedding from panel d, where blue regions denote “cold spots” (Getis-Ord test, *q-*value < 0.05) – areas of the embedding where control cells are depleted for neighbors of the titled perturbation (*egr2b* above, *cdx4;cdx1a* below). **f**-**g** UMAP plots in which cells are colored by the expression of individual DEGs (*epha4a, hoxb3a, hoxc3a,* or *hoxc6b; q*-value < 0.001) in controls or either *egr2b* or *cdx4;cdx1a* crispant neural progenitor cells .

**Figure 4.**
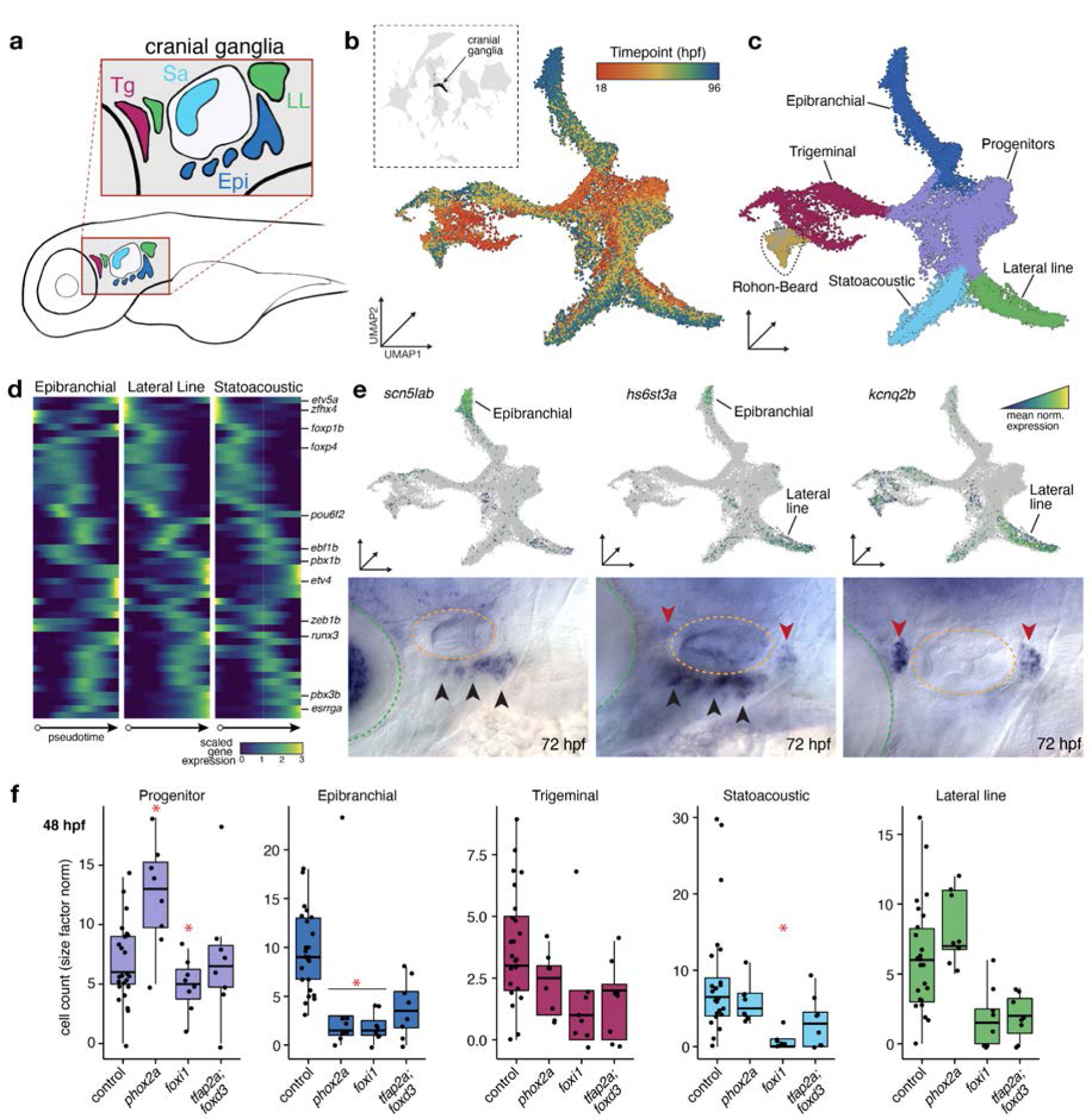
Whole embryo phenotyping robustly captures effects in the lowly abundant cells of the cranial sensory ganglia. **a,** A cartoon depiction of a lateral view of the sensory cranial ganglia in a ∼48 hpf zebrafish embryo. Colors represent ganglia types: Tg, Trigeminal ganglion; LL, lateral line ganglion; Epi, epibranchial ganglion; Sa, statoacoustic ganglion. **b-c**, Global UMAP embedding with cranial ganglia (n = 29,782) and Rohon-Beard neurons in black (inset). Sub-UMAP of cranial ganglia colored by time point (**b**) or cell type (**c**). These embeddings include both wild-type cells and cells from perturbation experiments. **d**, Row scaled pseudotime heatmaps of transcription factors enriched in one sensory ganglion trajectory branch. Genes listed on the y-axis are those with previously identified roles in cranial ganglia development. **e**, UMAP expression plots (above) and whole mount *in situ hybridization* (ISH) of embryos 72hpf (below) for three genes that are specific to either the epibranchial ganglion (left), lateral line ganglion (right), or both (middle) but are not expressed in other cells of the cranial sensory ganglia. Imaging of the lateral and anterior aspect, with eyes (light blue) and ears (yellow) are marked by dotted lines, and with arrowheads indicating epibranchial ganglia (black) or lateral line ganglia (red). **f**, Boxplots of the sensory cranial ganglia cell type counts from individual embryos at 48 hpf. *phox2a*, *foxi* and *tfap2a;foxd3* crispants. Significance is relative to wildtype control-injected embryos (* indicates *q-*value < 0.05).

Given the continuity of many of our trajectories from one cell type to another, we sought to understand the lineal relationships reflected in our data, e.g. the differentiation of mesodermal progenitors to myoblasts, to fast muscle myocytes (**Extended Data Fig. 5a**). However, inferring lineage relationships from transcriptional similarity alone can be fraught ^5, 19^, inspection of the UMAP embedding for the muscle trajectory suggested that slow and fast muscle cells share a common progenitor, however, it is known that slow muscle arises independently of fast muscle from precursors present prior to our earliest time point (18hpf) (**Extended Data Fig. 5b**). To distinguish between bona fide lineage relationships and mere transcriptional similarity between cell types across our atlas, we manually constructed a graph of documented lineage relationships, harmonized with our cell type annotations (**Extended Data Fig. 5c**).

**Figure 5.**
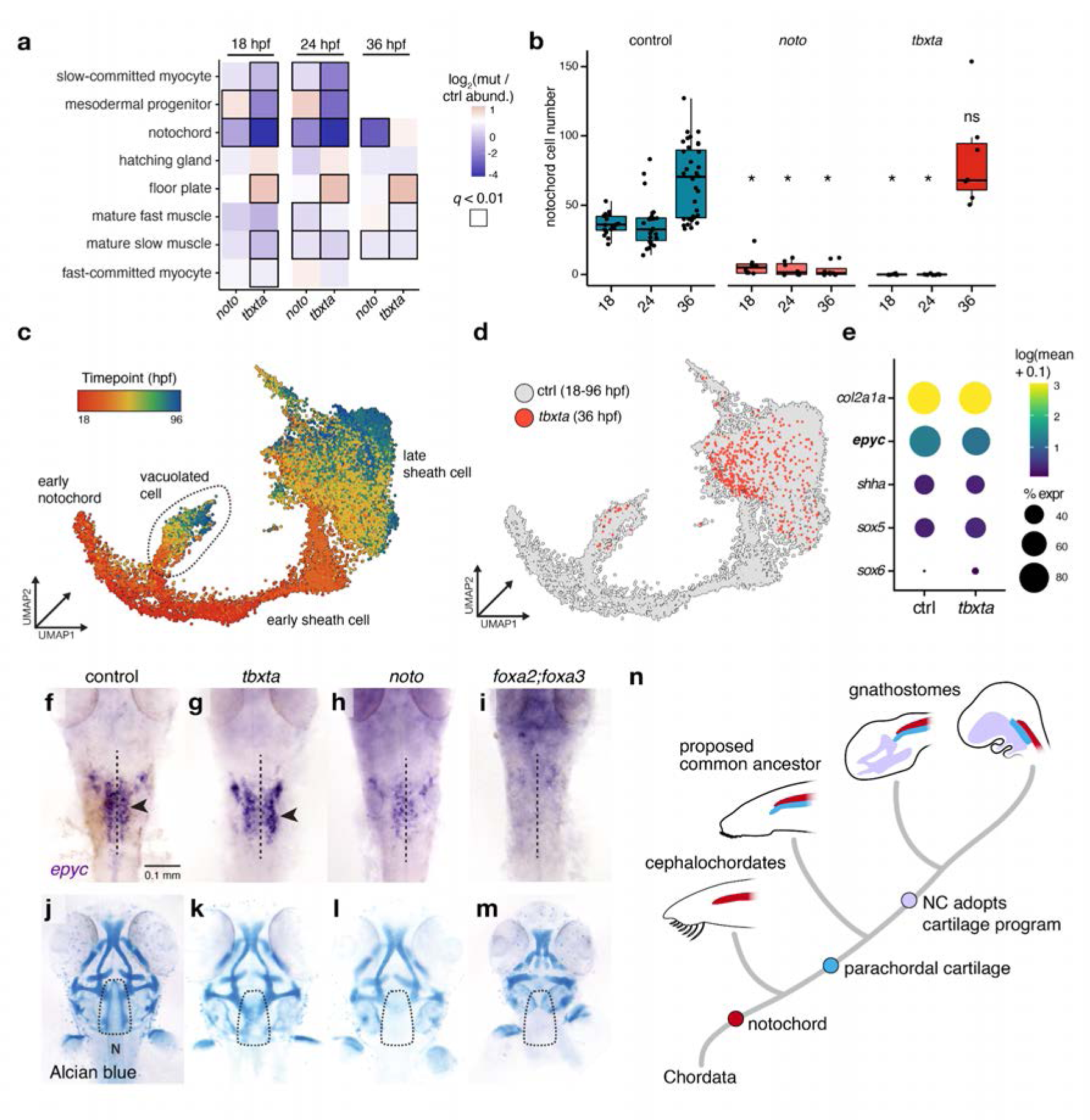
Tbxta and Noto perturbations uncover the genetic requirements of cranial cartilage development. **a**, Axial and paraxial mesodermal derivatives and their cell abundances relative to control embryos at three timepoints for *tbxta* and *noto* crispants. Black squares indicate significance (Beta-binomial regression, *q*-value < 0.01). **b**, Boxplots of notochord cell counts from individual embryos for controls, *noto* and *tbxta* crispants. Significance (* indicates q < 1e-5) is relative to wildtype control-injected embryos. **c**, UMAP embedding of the notochord trajectory constructed with reference cells and *tbxta* cells. Cells are colored by time point and are labeled by sub-type annotation. **d**, UMAP embedding of notochord cells, colored by genotype. **e**, A dotplot for a subset of genes that are expressed in notochord and *tbxta-*independent cells (referred to as NLCs). Color represents mean normalized gene expression, and circle size indicates the percent of notochord cells expressing the gene at 36 hpf. **f-i**, *epyc in-situ* hybridization (36 hpf; dorsal, anterior view) in control, *tbxta*, *noto*, and *foxa;foxa3* crispants. The dashed line indicates the notochord and parachordal cartilage cells in control and *tbxta* are marked by black arrowheads. **j-m**, Alcian blue staining of 72 hpf control, *tbxta*, *noto*, and *foxa2/foxa3* crispants. Dashed outline surrounds the parachordal cartilage region. All *tbxta*, *noto*, and *foxa2/foxa3* crispants lack a notochord. (N, notochord; dotted line surrounds the PC). **n**, A model depicting the hypothesized relationship between the notochord and cranial cartilage and bone elements over chordate evolution.

Using our individual-resolved, whole-organism data, we were also able to estimate the variability of cell type abundances over developmental time. To estimate variance, we adapted a statistical framework commonly used to account for mean-variance relationships in sequencing experiments to model variability in cell abundances ^20^. We found that most cell types vary in line with expectation given the nature of cell count data, but we did see excess variance in some cell types. Cell types that were significantly variable (*p*-value < 0.05; **Methods**) include the enveloping layer (EVL), mesodermal progenitor cells, and notochord cells at 20 hpf, and neural progenitor, optic cup, notochord, and head mesenchyme cells at 36 hpf (**Fig. 1e, Extended Data Fig. 6**). In addition to offering clues about the dynamic and transient nature of particular cell types, these variance estimates serve as important bases for our statistical assessment of perturbation-induced cell type abundance changes.

### Systematic genetic perturbation of embryonic development

Next, we used sci-Plex to label and measure single cell profiles across time from developing zebrafish F0 knockouts (crispants) generated by CRISPR-Cas9 mutagenesis (**Methods**). We first compared individual crispants with mutants deficient for *tbx16* or both *tbx16* and *msgn1*, which have well-studied phenotypes at 24 hpf^21^. Nearly all crispants were indistinguishable from stage-matched null mutants by gross morphology, displaying disorganized tail somite formation and the characteristic enlarged tail bud. We also looked for molecular or cellular differences between cells from knock-out (crispant or null) to controls. As previously documented ^22–24^, both exhibited a marked loss of slow and fast muscle and accumulated mesodermal progenitor cells, demonstrating the ability of our methodology to accurately pair genetic changes to loss of specific cell types (**Extended Data Fig. 7**).

We then scaled up our approach to profile many different genetic perturbations spanning multiple time points during embryogenesis (**Fig. 2a**). In total, we targeted 23 genes or gene pairs involved in the development of either mesoderm (*cdx4, cdx1a, tbxta, tbx16, tbx16l, msgn1, wnt3a, wnt8a, noto, smo, tbx1, hand2*), central or peripheral nervous system (*egr2b, epha4a, hoxb1a, mafba, zc4h2, phox2a, foxi1*, *hgfa, met*), or neural crest lineages (*foxd3, tfap2a*) (**Supplementary Table 3**). We designed 2-3 gRNAs per gene and checked for editing efficiency at target regions via a sequencing-based assay (**Extended Data Fig. 8; Supplementary Table 4**). A final set of gRNAs were chosen based on their ability to produce expected phenotypes in F0 knockouts without inducing non-specific cell death (**Extended Data Fig. 9**). For each gene target, we collected 8 embryos at an average of three of five timepoints that overlapped with the reference dataset: 18, 24, 36, 48, and 72 hpf. Altogether we profiled cells from 804 uniquely barcoded embryos across 98 conditions (including injection controls (n = 159), perturbations (n = 645), and multiple timepoints) and sequenced 2.7 million cells from a single sci-RNA-seq3 experiment and up to an estimated 10% of cells from each embryo (**Fig 2a**, **Extended Data Fig. 10**). Of these, the ∼600,000 cells from control-injected embryos did not display batch effects when co-embedded with our wild type time-series reference, and they are included in the final reference dataset (**Extended Data Fig. 3**).

To annotate cells by type for perturbed embryos and to facilitate cell type abundance analyses, we first projected the mutant data onto our reference atlas ^25^ and then transferred annotations using a fast, approximate nearest-neighbor algorithm (**Methods, Extended Data Fig. 11**). To assess perturbation-dependent cell type abundance changes, we transformed the data from a gene expression matrix into a cell type abundance matrix, effectively summarizing the number of each cell type observed within each embryo (**Fig. 2b**). After normalizing for the total cells recovered from each embryo, we performed dimensionality reduction to visualize this compositional data. Across the whole experiment, the primary source of variation in cell type proportions is embryo age (**Extended Data Fig. 12**). Within individual time points, perturbations with similar gross phenotypes readily grouped together – for example, loss of function for *tbxta* or *wnt3a;wnt8a*, all of which are important for maintenance of neuromesodermal progenitor cells (NMPs)^26^. In contrast, knocking out the hedgehog receptor *smoothened (smo)* resulted in a distinct cell type composition at the whole-embryo scale, consistent with the widespread requirements of hedgehog signaling during development^27^ (**Fig. 2c**).

### Using cell type composition as a phenotypic readout of genetic perturbations

To systematically discern and rank all changes in cell type abundances across perturbations, we applied a beta binomial regression model, which is well suited for assessing proportional changes in cell count data^28^ (**Methods**). To robustly measure changes in cell type abundance, we collected replicate embryos (n=8) for each perturbation/time point combination and compared them to stage-matched, control-injected embryos. Our analyses identified a range of significant differentially abundant cell types (DACTs) across the perturbations tested (**Fig. 2d, Extended Data Fig. 13**). For example, crispant embryos for transcription factors that regulate the development of early somitic lineages – Tbx16, Msgn1, and Tbx16l^22–24^ – exhibited both pronounced and subtle cell type abundance changes that were concordant between embryos (**Extended Data Fig. 14**). This suite of transcription factors regulates differentiation of the NMP population that gives rise to mesodermal progenitor cells (MPCs), and posterior spinal cord progenitors (pSCPs) (**Fig. 2e**)^21^. Accumulation of stalled MPCs has been well characterized in *tbx16/msgn1* single and double mutants; however, the consequences to the pSCP lineage have not been examined. Our data show that within individual embryos, both MPC and pSCP lineages are halted at the precursor stage in a graded fashion across single and double crispants (**Fig. 2f**). Thus, by examining whole transcriptomes, our data suggests that Tbx16, Tbx16l and Msgn1 interact to cooperatively control the differentiation of both mesodermal and neural progenitor cells from the NMP population and uncovers putative sets of new target genes for these transcription factors in both cell populations.

### Evaluation of differential gene expression across perturbations

To identify the transcriptional responses of each cell type to genetic perturbation, we performed differential gene expression (DE) tests to complement the differential abundance analysis. For each embryo, we combined cell data by type prior to testing (**Methods**). Pairwise DE tests between pseudo-bulked control and perturbed cells revealed an average of 1,470 differentially expressed genes (DEGs) for each perturbation, summed across all cell types (**Extended Data Fig. 15a**). Hierarchical clustering of DEGs highlighted that perturbations within a given genetic circuit induced common patterns of differential expression.

For example, we identified DEGs for neural progenitors for a suite of crispant perturbations that are known to affect neurogenesis (*cdx4, cdx1, wnt3a, wnt8a, mafba, hoxb1a, egr2b, smo,* and *epha4a*) (**Fig. 3a**). While these perturbations did not result in robust cell type composition changes, we nevertheless uncovered many perturbation-induced DEGs (**Extended Data Fig. 16**). Knocking out genes important for hindbrain development – *egr2b*, *mafba, epha4a,* and *hoxb1a*^29–31^ – exemplified this phenomenon (**Fig. 3b**). Prior studies have revealed important roles for these factors in the segmentation and specification of neural progenitor cells in the hindbrain, but cell type-specific transcriptome-wide consequences of loss-of-function are unknown. When we compared DEGs for these perturbations, they form two major groups in accordance with known genetic interactions^32–34^.

Moreover, DEGs are enriched for biological processes and pathways involved in brain and nervous system development, offering new hypotheses for gene effectors of our target genes (**Fig. 3c**).

Because neural progenitor cells at these stages have generally similar transcriptional programs and do not form distinct boundaries in low dimensional space, we additionally sought to identify perturbation-dependent shifts in transcriptional states that were cluster agnostic (**Fig. 3d**). We used the Getis-Ord test to identify regions of the reference UMAP embedding that were either enriched or depleted of perturbed cells in a co-embedded subset of the data (**Methods**)^35^. This analysis revealed distinct regions of the reference UMAP space that were depleted for perturbed hindbrain neural progenitor cells (**Fig. 3e, Extended Data Fig. 15b)**. These regions corresponded with differential gene expression, such as a significant downregulation of *epha4a* expression in *egr2b* crispant neural progenitors, consistent with previous work^36^ (**Fig. 3f**). Prior studies of *cdx1* and *cdx4* identified functions during posterior mesoderm development, where they coordinate multiple pathways and activate hox gene expression^37–40^. Studies of zebrafish *cdx4;cdx1a* mutants also revealed the importance of these genes in hindbrain patterning^41^. Indeed, we find that three hox genes are significantly downregulated in *cdx4* and *cdx4;cdx1a* crispant neural progenitor cells (**Fig. 3g**). More broadly, our whole-embryo, single cell measurements across time now enable a comprehensive view of candidate targets for these key transcription factors. These analyses highlight our ability to leverage individual-level transcriptome measurements to systematically evaluate perturbation-dependent transcriptional changes in each cell type and provide new hypotheses for functional studies.

### Dissecting the molecular identities of the cranial sensory ganglia

Specialized subsets of some cell types can express highly similar transcriptomes despite having distinct functions, lineage origins, or anatomic locations ^42–44^. Alternatively, cell types arising from distinct lineal origins can give rise to identically functioning cells ^5, 8, 45^. Disentangling these unique scenarios may not be possible from snapshots of normal development, regardless of the resolution of the data. The cranial sensory neurons, which transmit information from the head, ear, heart and viscera, are examples of a cell type that has been difficult to study in zebrafish owing to their relatively low cellular abundance in the embryo, complex developmental history, and a lack of known markers to distinguish their subtypes ^46^. Despite their scarcity, we captured ∼30,000 cranial sensory neurons (∼20 cells/embryo) contained within a single cluster, which formed four distinct branches upon sub-clustering. To identify whether these branches reflected placodal origins, neuronal function, or something else, we manually compared branch-specific gene expression with published expression data. We hypothesized that consistent with their distinct placodal origins, the branches represent the epibranchial, trigeminal, statoacoustic, and lateral line cells, all radiating from a putative set of progenitors (**Fig. 4a-c; Extended Data Fig. 17a**).

We next sought to characterize the molecular differences between the subtypes of cranial sensory neurons and to identify the putative lineage determining factors that distinguish them. Differential expression analysis identified 45 transcription factors that were expressed in the progenitors and just one of the daughter branches (**Fig. 4d**). This set of genes included some factors identified to regulate sensory neuron development^47^, but the majority have no previously reported role for these ganglia. To validate our cell type annotations and characterize novel subtype markers, we additionally selected 11 terminally-expressed genes to analyze by whole mount in situ hybridization (ISH). We were able to synthesize *in situ* hybridization probes for 9 of these and found 8 that labeled the expected sensory ganglia at 72 hpf, establishing a new set of molecular markers for these subpopulations (**Fig. 4e; Extended Data Fig. 17d**).

To explore the genetic requirements of the cranial sensory ganglia, we disrupted two transcription factors that are important for their development: *foxi1* and *phox2a*^48–52^. *Foxi1* is expressed early in development in placodal progenitor cells and is required broadly for proper differentiation of cranial ganglia neurons. *Phox2a* is required downstream of *foxi1* for development of epibranchial neurons, where it is specifically and robustly expressed (**Extended Data Fig. 17b**). Consistent with prior studies, we found that loss of *phox2a* led to a significant reduction of epibranchial neurons and an increase in progenitor cells, suggesting that these cells have stalled in a progenitor state. In *foxi1* crispants, progenitor cells and all four classes of cranial sensory ganglia were reduced, consistent with the early requirement of *foxi1* in placodal precursors of these lineages (**Fig. 4f, Extended Data Fig. 17c**).

Although cranial sensory ganglia are known to have both ectodermal placode and neural crest origins across different species ^53–55^, the lineage contributions to each of the cranial ganglia in zebrafish are unclear. Zebrafish cranial ganglia arise early in development predominantly from ectodermal placodes; later on, the neural crest contributes to trigeminal ganglia and potentially other classes^56, 57^. In *tfap2a;foxd3* crispants, for which corresponding mutants lack all neural crest derivatives^58^, we predicted that if neural crest cells contributed to specific ganglia, that we would detect corresponding decreases in cell abundance. We identified mean reductions (50-70%) in numbers of neurons of the trigeminal, epibranchial, statoacoustic, and lateral line ganglia but not progenitors at 48 hpf (**Fig. 4f**).

Moreover, epibranchial and lateral line ganglia cells were consistently reduced across all three timepoints collected (36, 48, and 72hpf). While we were underpowered to assign significance to these reductions, the data suggest that neural crest contributions to zebrafish cranial ganglia are likely more widespread than previously appreciated. Precise mapping of the genetic architecture of rare cell types (like the sensory ganglia) will require either an alternative experimental design, more replicates, or technological advances (e.g. increased cell capture rate). Nevertheless, these experiments demonstrate that deep single-cell transcriptome analysis of whole wild type and mutant embryos can reveal distinguishing molecular features, genetic requirements and lineage contributions of cells comprising a small percentage (less than 1%) of total cells within the embryo.

### Single cell sequencing reveals a shared transcriptional program for notochord and cranial cartilage

Because the notochord is the defining feature of chordates and serves critical structural and signaling roles in the vertebrate embryo ^59^, we targeted two highly conserved transcription factors essential for its development: *noto* and *tbxta*/*brachyury* ^59–62^. Our differential cell type abundance analyses largely reflected the expected phenotypes for *noto* and *tbxta*, e.g. reduced slow muscle and notochord cells, and increased floorplate cells in *tbxta* crispants ^61^ (**Fig. 5a**). In both *noto* and *tbxta* crispants, there is a near complete loss of notochord cells at both 18 and 24 hpf. However, despite the absence of a visible notochord, we detected a near complete recovery of putative notochord cells by 36 hpf in *tbxta* crispants (**Fig. 5b**).

To investigate these unexpected cells (referred to as NLCs – notochord-like cells), we refined our annotations to distinguish the developmental trajectories of the two cell types that comprise the notochord: inner vacuolated cells and outer sheath cells (**Fig. 5c**). Vacuolated cells aid in embryonic axis elongation, while sheath cells form a surrounding epithelial layer that secretes a collagen-rich extracellular matrix around the notochord ^63, 64^. In *tbxta* crispants a majority of NLCs resembled maturing wild type sheath cells (**Fig. 5d**). Comparison of NLCs relative to wild type sheath cells revealed 157 differentially expressed genes, but only a handful uniquely marked NLCs (*q-*value < 0.01, **Extended Data Fig. 18b**). At this point our mutant data had unmasked NLCs, a cryptic, sheath cell like cell type (*epyc+*, *col2a1a+, shha+*) (**Fig. 5e**), arising between 24 and 36hpf, despite the absence of a visible notochord.

To anatomically locate NLCs, we visualized the spatial localization of *epyc* expression using whole mount ISH. In control embryos, *epyc* is expressed weakly throughout the notochord and strongly in parachordal cartilage (PC), a conserved, mesodermally-derived cartilage structure that later develops into the cranial base of the skull (**Fig. 5f**)^65, 66^. Furthermore, a putative NLCs marker revealed by our differential analysis, *tgm2l*, labeled PC cells but not notochord in wild-type embryos (**Extended Data Fig. 18c**). Consistent with the proposed similarities of the notochord sheath to cartilage ^59^, we found that both cell types share the core conserved module of gene expression for cartilage formation (*sox5/6/9*, *col2a1a*) (**Fig. 5e**). Thus, the apparent and unexpected “recovery” of notochord cells in *tbxta* crispants revealed that the NLCs, which are transcriptionally nearly indistinguishable from notochord sheath cells, are indeed parachordal cartilage (PC) cells.

The similarity between PC and notochord led us to wonder how their genetic requirements overlapped, so we visualized these cells in embryos lacking the lineage determining factors *noto* and *tbxta*. In *tbxta* crispants at 36 hpf, while notochord cells are missing, *epyc*+ early PC cells are present (**Fig. 5f,g**). In *noto* mutants, *epyc* is weakly expressed by some cells in the posterior head, but these cells lack any organization around the midline. We next determined whether the *tbxta*-independent, early PC cells retained the ability to mature into chondrocytes by staining head cartilage at 72 hpf (**Fig. 5j-l, Extended Data Fig. 19**). While Alcian-positive PC cells are present at 72 hpf in control and *tbxta* crispant embryos, posterior PC does not form in *noto*, consistent with the lack of *epyc*+ precursor cells (**Fig. 5h**). Thus, *tbxta* and *noto* have separate functions during PC and notochord development.

To probe the earlier genetic requirements of these cells, we generated crispants for both *foxa2* and *foxa3*, two TFs with conserved roles during axial mesoderm specification. In mice, *foxa2* alone is required for notochord development, whereas in zebrafish, knockdown of *foxa2* and *foxa3* together leads to loss of all axial mesoderm derivatives ^67, 68^. We found that in the absence of both *foxa2* and *foxa3*, the notochord fails to develop, *epyc*+ PC cells are missing, and no PC forms by 72 hpf (**Fig. 5i,m, Extended Data Fig. 20a,b**). Thus, while both the notochord and PC derive from the early embryonic *foxa2/3*-dependent axial mesoderm progenitor pool ^69, 70^, notochord development additionally requires *noto*, and *tbxta*, whereas PC development only requires *noto*. Overall, this result suggests that PC and notochord fate divergence occurs early in the axial mesoderm and that genetic requirements in PC differ from other axial tissues (**Extended Data Fig. 20c**).

The high degree of transcriptional similarity and differing genetic requirements of parachordal cartilage cells (“true cartilage”) and notochord sheath cells (“cartilage-like”)^71^ offers clues into the evolutionary origin of vertebrate cranial skeletons. While it is now clear that much of the anterior head cartilage is neural crest-derived, the evolutionary origin of the ancient mesodermal head cartilage, which produces the posterior skull, is unknown ^65, 66^. Based on the shared location, gene expression, and transcriptional regulation of the progenitors for PC and notochord, we speculate that the cartilage-like notochord cells are the direct precursors to skeletal cranial elements in the vertebrate lineage. Thus, we suggest that as creatures evolved from an amphioxus-like vertebrate ancestor, some of the embryonic anterior notochord cells split to form the PC just lateral to the notochord, which allowed the development of more complex mesodermal cartilage structures. Later, these joined with neural crest-derived cartilage to form the modern vertebrate skull (**Fig. 5h**)^72^. These findings highlight the promise of high-resolution molecular phenotyping to deepen our understanding of the relationship between gene expression and genetic networks, facilitating new hypotheses about the evolutionary origins of individual cell types.

## Discussion

Here, we present a new approach (whole-organism labeling) and dataset for systematically analyzing the impact of genetic perturbations on each cell type in the developing zebrafish at single-cell resolution. To achieve this, we first established an individual-resolved reference atlas of zebrafish development. Our data fills a gap in existing zebrafish atlas datasets ^7, 8, 17^, provisioning a single cell dataset spanning 18 to 48 hours post fertilization in 2 hour increments. This developmental period is epitomized by the differentiation of diverse cell types throughout the organism, and the accompanying cell type annotations reflect this richness (33 major tissues, 85 broad cell types, and 156 cell subtypes). All subsequent perturbation experiments leverage this reference data to rapidly annotate and assess the cellular consequences of diverse genetic mutations.

We profiled 23 genetic perturbations, collecting 2.7 million single cell transcriptomes, from 804 embryos across 98 conditions in a single sequencing experiment. The degree of replication (8 or more embryos per perturbationxtimepoint) is unprecedented in single cell experiments and affords the statistical power to comprehensively detect gains and losses in the abundance of both common and rare cell types throughout the embryo. For example, dissected the molecular signatures of four cranial ganglia neuron types and their precursors, which together comprise less than 1% of the cells in the organism. Additionally, whole-organism profiling alleviates the need for complex reporter systems or other means of purifying cells of interest *a priori*^73^. This feature of our approach is highlighted by the comparison of *tbxta* and *noto* mutants, both of which fail to develop notochords. Further investigation of these mutants identified the parachordal cartilage as transcriptionally indistinguishable from notochord sheath cells and revealed independent genetic requirements for these two cell types, a finding that provides new clues about the origins of the vertebrate skull.

Our method is not without limitations for future research to address. First, while we are well-powered to detect changes in certain lowly abundant cell types, the statistical power required is still dependent on the magnitude of the effect. Additionally, while observing phenotypes in a whole organism context offers advantages, it quickly becomes infeasible to profile larger organisms that may contain billions to trillions of cells. Finally, while we assessed mutagenesis efficiency at the whole-embryo level prior to single cell sequencing, low levels of mosaicism in F0 crispants is a concern, especially when this approach is used for morphogens or other secreted factors where a small amount of mosaicism may be sufficient to rescue a mutant phenotype. An ideal assay would capture both the single cell transcriptome, and the perturbed genetic allele, allowing for the interpretation of perturbations with no apparent phenotype.

Looking forward, we anticipate that the ease of this framework – collection of mutant transcriptomes coupled with rapid annotation and cellular and molecular comparison to a reference – will lead to the development of a cumulative, community resource. Moreover, the data collected in the future is likely to grow, both quantitatively (timepoints, mutants, environmental perturbations, and species) (see accompanying manuscript, Dorrity *et al.* 2022), but also qualitatively (in richness). For example, the combination of single embryo perturbation studies with lineage tracing, with the co-measurement of the epigenome with the transcriptome, or with spatial readouts of morphology, will afford new avenues to interrogate the atlas and deepen our understanding of development. The resulting integrated resource is also likely to spur new generalized machine learning models of gene regulation, akin to those that have revolutionized the fields of protein design and natural language processing ^74, 75^.

Together, these technical and analytical advances will facilitate a new era of developmental genomics that moves towards a comprehensive, multi-scale understanding of embryogenesis.

## Methods

### Animal rearing, staging and stocks

Staging followed ^79^ and fish were maintained at ∼28.5°C under 14:10 light:dark cycles. Fish stocks used were: wild-type AB, noto^n1^,_60_ tbx16^b104,22^ Tg(isl1:gfp)^rw0^, Tg(p2rx3:gfp)^sl1^, mafba^b337^,^30^ hgfa^fh528^, met^fh533^ ^80^. Fish were anesthetized prior to imaging or dissociation with MS222 and euthanized by overdose of MS222. All procedures involving live animals followed federal, state and local guidelines for humane treatment and protocols approved by Institutional Animal Care and Use Committees of the University of Washington and the Fred Hutchinson Cancer Research Center.

### Imaging

For confocal imaging (**Extended Data Fig. 9**) embryos were anesthetized with MS222 and mounted in 1% low-melt agarose on a coverslip and imaged on an LSM700 inverted confocal microscope at 20x magnification. Colorimetric RNA in situs were cleared in glycerol, dissected to remove yolk granules, flat-mounted between two coverslips separated by high vacuum grease posts, and imaged on a Zeiss Axioplan transmitted light microscope.

### CRISPR-Cas9 mutagenesis in zebrafish embryos

Guide RNAs were designed using either the Integrated DNA Technologies (IDT) or CRISPOR^81^ online tools. Guide RNA and RNP preparation closely follow a recently published protocol for efficient CRISPR-Cas9 mutagenesis in zebrafish ^15^. Briefly, guide RNAs were synthesized as crRNAs (IDT), and a 50µM crRNA:tracrRNA duplex was generated by mixing equal parts of 100µM stocks. Cas9 protein (Alt-R S.p. Cas9 nuclease, v.3, IDT) was diluted to a 25µM stock solution in 20nM HEPES-NaOH (pH 7.5), 350mM KCl, 20% glycerol. The RNP complex mixture was prepared fresh for each injection by combining 1µL 25µM crRNA:tracrRNA duplex (with equal parts each gRNA per gene target), 1µL 25µM Cas9 Protein, and 3µL nuclease-free water. Prior to injection, the RNP complex solution incubated for 5 min at 37℃ and then kept at room temperature. Approximately 1-2nL was injected into the cytoplasm of one-cell stage embryos.

### Genotyping

Two days after CRISPR-Cas9 RNP injections (48 hpf), pools of five F0-injected embryos for each gRNA set were lysed in 100μl alkaline lysis buffer (25mM NaOH, 0.2mM EDTA) and heated at 95 °C for 30 min. The solution was neutralized by an equal volume of neutralization buffer (40 mM Tris-HCl, pH 5.0). Rhamp-seq primers were designed using the Rhamp-seq IDT design tool. Rhamp-seq primers were reconstituted in low-TE buffer (10mM Tris/HCl ph 7.4, 0.1mM EDTA) to a final concentration of 10μM. These primers were then mixed in 4 pools as specified by the IDT design tool (Pool1-FWD, Pool1-REV, Pool2-FWD, and Pool2-REV). Each primer in these pools was mixed such that the primer’s final concentration in the pool was 0.25μM. Genotyping PCRs for each crispant were performed using 5μL of 4x Rhampseq Master Mix 1 (IDT), 2μL of FWD pool, 2μL of REV pool, and 11μL of gDNA template. 20 cycles of PCR were performed using the following thermocycler program:

1. 95°C for 10 minutes
2. 95°C for 15 seconds
3. 61°C for 4 minutes
4. Return to step 2 for 10 cycles total
5. 99.5°C for 15 minutes

Following amplification, PCR products were purified using a 1.5X SPRI bead cleanup (Beckman Coulter, cat. no. A63880) and eluted in 15μL low-TE buffer. Index PCR was performed using 5μL of 4x Rhampseq Master Mix 2, 2μL of Indexing PCR primer (i5), 2μL of Indexing PCR primer (i7), and 11μL of purified PCR product. An additional 20 cycles of index PCR were then performed using the following thermocycler program:

1. 95°C for 10 minutes
2. 95°C for 15 seconds
3. 60°C for 30 seconds
4. 72°C for 30 seconds
5. Return to step 2 for 20 cycles total
6. 72°C for 1 minutes.

After the index PCR, sequencing libraries were pooled, purified with a 1x SPRI bead cleanup and sequenced on the Illumina MiSeq 600 cycle kit with 2x300 cycle paired-end reads. Reads were analyzed using the ampliCan software package with default settings and standard vignette workflow^77^.

### Immunohistochemistry and labeling

Alkaline phosphatase *in situ* hybridization was performed using standard conditions^82^. Alcian blue staining followed an online procedure (SDB Online Short Course, Zebrafish Alcian Blue), except that embryos were raised in 1-phenyl-2-thiourea (Millipore-Sigma, cat. no. P7629) to suppress pigment formation rather than bleaching.

### Preparation of barcoded nuclei

Individual zebrafish embryos (18 to 96 hpf) were manually dechorionated with forceps and transferred to a 10cm petri dish containing 1X TrypLE (Thermo Fisher, cat. no. 12604013). Using a wide bore tip, embryos were transferred, one by one, into separate wells of a 96-well V-bottom plate containing 75μL of 1X TrypLE (Thermo Fisher, cat. no. 12604013) + 2 mg/mL Collagenase P (Millipore Sigma, cat. no. 11213865001). Embryos were then dissociated by 10 strokes of manual trituration at 30℃ once every 5 minutes. Dissociation continued until no visible chunks were present under a dissecting scope, which took between 20-40 minutes depending on embryo stage (e.g. 20 minutes for 18hpf and 40 minutes for 72 hpf). Stop solution [1x dPBS (Thermo Fisher cat. no. 10010023), 5% FBS (Thermo Fisher cat. no. A4736401)] was then added to each well to quench the proteases. Cells were then spun down at 600xg for 5 min. Cells were then resuspended in 200μL in cold dPBS and spun down again.

After rinsing, the supernatant was removed fully and cells were resuspended in 50μL of CLB [Cold Lysis Buffer – 10mM Tris/HCl pH 7.4, 10mM NaCl, 3mM MgCl_2_, 0.1% IGEPAL, 1% (v/v) SuperaseIn RNase Inhibitor (20 U/µL, Ambion), 1% (v/v) BSA (20 mg/ml, NEB)] + 5μL of hash oligo (10 uM, IDT) and incubated for 3 min on ice. Following lysis, 200μL of ice cold, 5% fixation buffer [5% paraformaldehyde (EMS, cat. no. 50-980-493), 1.25x dPBS] was added to each well. After an additional round of mixing, nuclei were fixed on ice for 15 minutes. All wells were then pooled together in a 15 mL conical tube and spun down for 15 min at 750xg. Supernatant was decanted and cells rinsed in 2mL of cold NBB [Nuclei Buffer + BSA – 10mM Tris/HCl pH 7.4, 10mM NaCl, 3mM MgCl_2_, 1% (v/v) BSA, 1% (v/v) SuperaseIn RNase Inhibitor] at 750xg for 6 min. Supernatant was then carefully aspirated and the nuclei were resuspended in 1mL of NBB and flash frozen in LN_2_ and stored at -80°C.

### sci-RNA-seq3 library construction

The fixed nuclei were processed similarly to the published sci-RNA-seq3 protocol^6^ with some modifications. Briefly, frozen, paraformaldehyde-fixed nuclei were thawed, centrifuged at 750xg for 6 min, and incubated with 500μL NBB (see previous) including 0.2% (v/v) Triton X-100 for 3 min on ice. Cells were pelleted and resuspended in 400μL NBB. The cell suspension was sonicated on low speed for 12s (Diagenode, Bioruptor Plus). Cells were then pelleted at 750xg for 5 min prior to resuspension in NB + dNTPs. The subsequent steps were similar to the original sci-RNA-seq3 protocol (with paraformaldehyde fixed nuclei) with some modifications, and a detailed, step-by-step protocol will be available as a supplementary protocol.

### Sequencing, read processing, and cell filtering

Libraries were sequenced on either an Illumina NextSeq 500 (High Output 75 cycle kit), Nextseq 2000 (P2 100 cycle kit), or Novaseq 6000 (S4 200 cycle kit) with sequencing chemistries compatible with library construction and kit specifications. Standard chemistry: I1 - 10bp, I2 - 10bp, R1 - 28bp, R2 - remaining cycles (> 45). Read alignment and gene count matrix generation was performed using the Brotman Baty Institute (BBI) pipelines for sci-RNA-seq3 (https://github.com/bbi-lab/bbi-dmux; https://github.com/bbi-lab/bbi-sci). After the single cell gene count matrix was generated, cells with fewer than 200 UMIs were removed, followed by removal of cells with UMIs greater than 3 standard deviations from mean. For mitochondrial signatures, we aggregated all reads from the mitochondrial chromosome and cells with > 25% mitochondrial reads were removed. Each cell was assigned to a specific zebrafish embryo based on the enrichment of a single hash oligo, as described previously^13^. Enrichment cutoffs were set manually based on the distribution of enrichment ratios (**Supplementary Table 1**). Removing cells with low hash enrichment ratios removed most doublets ^13^. Additional clusters of doublets, not removed using this procedure were manually inspected for marker genes and removed.

### scRNA-seq analysis

After RNA and hash quality filtering, data was processed using the Monocle3 workflow defaults except where specified: *estimate_size_factors()*, *detect_genes(min_expr = 0.1)*, *preprocess_cds()* with 100 principal components (using all genes) for whole-embryo and 50 principle components for subsets, *align_cds(residual_model_formula_str = “∼log10(n.umi)”)*, *reduce_dimension(max_components = 3, preprocess_method = ’Aligned’)*, and finally *cluster_cells(resolution = 1e-4)*.

### Hierarchical annotation and sub-clustering

To build maps where cluster annotations corresponded broadly to cell types, we first split the global reference dataset into 4 major groups that each contained either the epidermis, muscle, CNS neurons, or mesenchyme cells, along with other nearby cell types. Each of these groups was re-processed, embedded in 3 dimensions with UMAP and sub-clustered. Cluster resolution was optimized such that major groups were composed of 30 to 70 clusters that qualitatively represented the transcriptional diversity in a given set. Clusters were then assigned annotations based on the expression of marker genes (**Supplementary Table 8**) based on literature by an unsupervised signature-scoring method using anatomical-term gene lists from the Zfin database. With the exception of a few additional sub-clustering examples (i.e. the cranial ganglia), each cluster was assigned on “cell_type_sub” annotation. These subtype annotations were manually merged into “cell_type_broad” classifications based on cluster proximity or cell type functional groupings. We further merged these annotations into “tissue” groups based on whether broad cell types together composed a broader tissue. Finally, we designated each cell type into a “germ_layer” group based on the known germ layer of origin.

### Individual-level composition analysis

After cell type annotation, counts per cell type were summarized per embryo to generate an embryo x cell type matrix. Embryo composition size factors were calculated independently for each time point. The embryo x cell type matrix was stored as a cell_data_set object, allowing for preprocessing (PCA) and dimensionality reduction (UMAP) using the standard Monocle3 workflow.

### Query dataset projection and label transfer

The PCA rotation matrix, batch correction linear model, and UMAP transformation were computed and saved during the processing of the reference dataset. This computation was done on two levels: first, with all combined reference cells (global reference space), and second, in each of four sub-groups (sub reference space).

The query dataset was first projected into the global reference space using the following procedure: The PCA rotation matrix, which contains the coefficients to transform gene expression values into PCA loadings, was applied to the query dataset. The batch correction model was then applied to the resulting query PCA matrix to remove the effects of UMI count. Finally, the reference-calculated UMAP transformation was applied to the batch-adjusted PCA loadings to project the query data into the stable reference coordinate space. This procedure is similar to the procedure used in Andreatta *et al*.^83^

One of four major sub-group labels was transferred (mesoderm, mesenchyme-fin, periderm, CNS) using the majority label of its annotated nearest neighbors (k=10). Nearest neighbors were calculated using annoy, a fast, approximate nearest-neighbor algorithm (https://github.com/spotify/annoy). The query dataset was split into four sub-groups based on these assigned major group labels. Each query sub-group was projected into the sub-reference spaces using the corresponding saved PCA, batch correction, and UMAP transformation models using the same projection procedure. Finer resolution annotations (germ layer, tissue, broad cell type, sub cell type) were transferred in this sub-space using the majority vote of reference neighbors (k=10).

### Differential expression testing

Prior to differential expression testing, expression values were aggregated for each embryo across each cell type into “pseudo-cells”. We pooled embryos across time points and only compared embryos from the same sets of timepoints in each test. Differential expression analysis for pseudo-cells was performed using generalized linear models, as described previously^13^, with modifications to account for differential underlying count distributions in the “fit_models()” function in Monocle3^6^.

### Spatial autocorrelation of transcriptional responses to perturbation

The local spatial statistic Getis-Ord index (G_i_) was used to identify statistically significant regions of the UMAP embedding that were enriched or depleted of perturbed cells. A high-value G_i_ indicates that a perturbed cell is surrounded by other cells with the same perturbation, whereas a G_i_ close to zero indicates that a perturbed cell is surrounded by cells with other perturbation labels. A G_i_ was calculated for each cell’s local neighborhood (k=15) using the “localG()” function in the spdep package. This returns a z-score that indicates whether the observed spatial clustering is more pronounced than expected by random. Multiple testing correction was done using a Bonferroni correction. Areas of the UMAP where a given perturbation is enriched are called “hot spots” while areas where a given perturbation is depleted are referred to as “cold spots”.

### Statistical assessment of cell abundance changes

Changes in the proportions of each cell type were assessed by first counting the number of each annotated cell type in each embryo. To control for technical differences in cell recovery across embryos, “size factors” by dividing the total number of cells recovered from an embryo by the geometric mean of total cell counts across all embryos. The number of cells of each type recovered from each embryo were then divided by that embryo’s size factor.

Normalized counts for each cell type at time were then compared across genotypes using a generalized linear model defined by the equations:

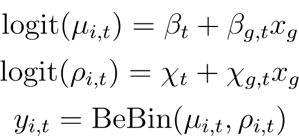

Where, the normalized counts of cell type at time is modeled as a beta-binomially distributed random variable with mean and “litter effect” (i.e. overdispersion with respect to the binomial distribution). We modeled both parameters of the beta binomial response as a function of genotype, reasoning that crispants might exhibit greater variability than wild-type embryos. The binary indicator variable denotes whether gene is knocked out in each embryo. Separate models for each gene in each cell type and at each time point were fit using the VGAM package ^84^. Significance of knockout effects in each model were assessed by Wald test on *β_g,t_* .

### Gene set enrichment analyses

After differential expression testing, genes that had significant coefficients (*q*-value < 0.05) were used for gene set enrichment analysis with the g:Profiler2 R package ^85^. Gene sets were filtered for significance (p < 0.05), and of the top gene sets, those having to do with neuronal development processes were chosen for visualization.

### Comparison of published zebrafish developmental atlases

Datasets for each study ^7, 8, 17^ were downloaded. The authors of each dataset had used different naming conventions for gene names. To harmonize the datasets, the gene names from each dataset were first converted to the GRCz11 ENSEMBL gene names. Genes with duplicated names were removed and only genes found in all three datasets were retained. Datasets were then aligned with the *IntegrateData* function in Seurat3. For **Extended Data Fig. 2c**, where wild type transcriptomes at 24 hpf are compared to stage matched transcriptomes from ^7, 8, 17^, wild type reference data was first downsampled, and then integrated using reciprocal PCA. Default hyperparameters were used for integration, PCA and dimensionality reduction. Following co-embedding, labels were transferred from ^7, 8, 17^ to the wild type reference data in the co-embedded space using the majority label from the 10 nearest neighbors. These labels were then used to calculate the concordance between the two datasets (**Extended Data Fig. 2d**).

## ACKNOWLEDGEMENTS

We thank Nola Klemfuss and the Brotman Baty Institute Advanced Technology Lab for support with sequencing and the data processing pipeline. We also thank Frank Steemers and Fan Zhang for additional sequencing support. We thank Ben Hamilton for custom illustrations. We thank T. Kaneko and J. Stonick for help with live imaging of sensory neurons and R. Garcia for assistance with fish husbandry and breedings.

## Author contributions

L.M.S., S.S., and C.T. conceived the project. L.M.S., D.K., C.B.M., D.R., and C.T designed experiments. L.M.S. and S.S. developed techniques and performed sci-RNA-seq3 experiments.

L.M.S. and D.K. performed all micro-injections. L.M.S., S.S. and M.W.D. did dissociation and nuclei collections. L.M.S. and S.S. performed computational analyses with M.D. and B.E. L.M.S., M.W.D., D.K., and C.B.M. annotated cell types in the developmental reference. D.K. performed the ISH. C.B.M., D.K., and L.M.S. performed imaging experiments. L.M.S., S.S., and C.T. wrote the manuscript with input from all coauthors. C.B.M., D.R., D.K., and J.S. contributed methods, supervision, and edited the manuscript. C.T. supervised the project.

## Data Availability

The datasets generated and analyzed during the current study are available in the NCBI Gene Expression Omnibus (GEO) repository: GSE202639.

## Code Availability

Pipelines for generating count matrices from sci-RNA-seq3 sequencing data are available at https://github.com/bbi-lab/bbi-dmux and https://github.com/bbi-lab/bbi-sci. Analyses of the single cell transcriptome data were performed using Monocle3, which was updated to include methods from this study; a general tutorial can be found at http://cole-trapnell-lab.github.io/monocle-release/monocle3. Additional custom analysis functions and scripts will be made available via a Github repository (https://github.com/cole-trapnell-lab/sdg-zfish).

## Supplementary Information

**Supplementary Table 1 -** Metrics from all scRNA-seq experiments

**Supplementary Table 2** - Hierarchical cell type annotations and evidence

**Supplementary Table 3** - List of perturbations and selected citations

**Supplementary Table 4** - gRNA sequences

**Supplementary Table 5** - rhAmp-Seq primer sequences

**Supplementary Table 6** - Metrics for perturbations

**Supplementary Table 7** - Cranial ganglia sub-cluster top marker genes

**Supplementary Table 8** - Top markers for each cell subtype

**Extended Data Figure 1.**
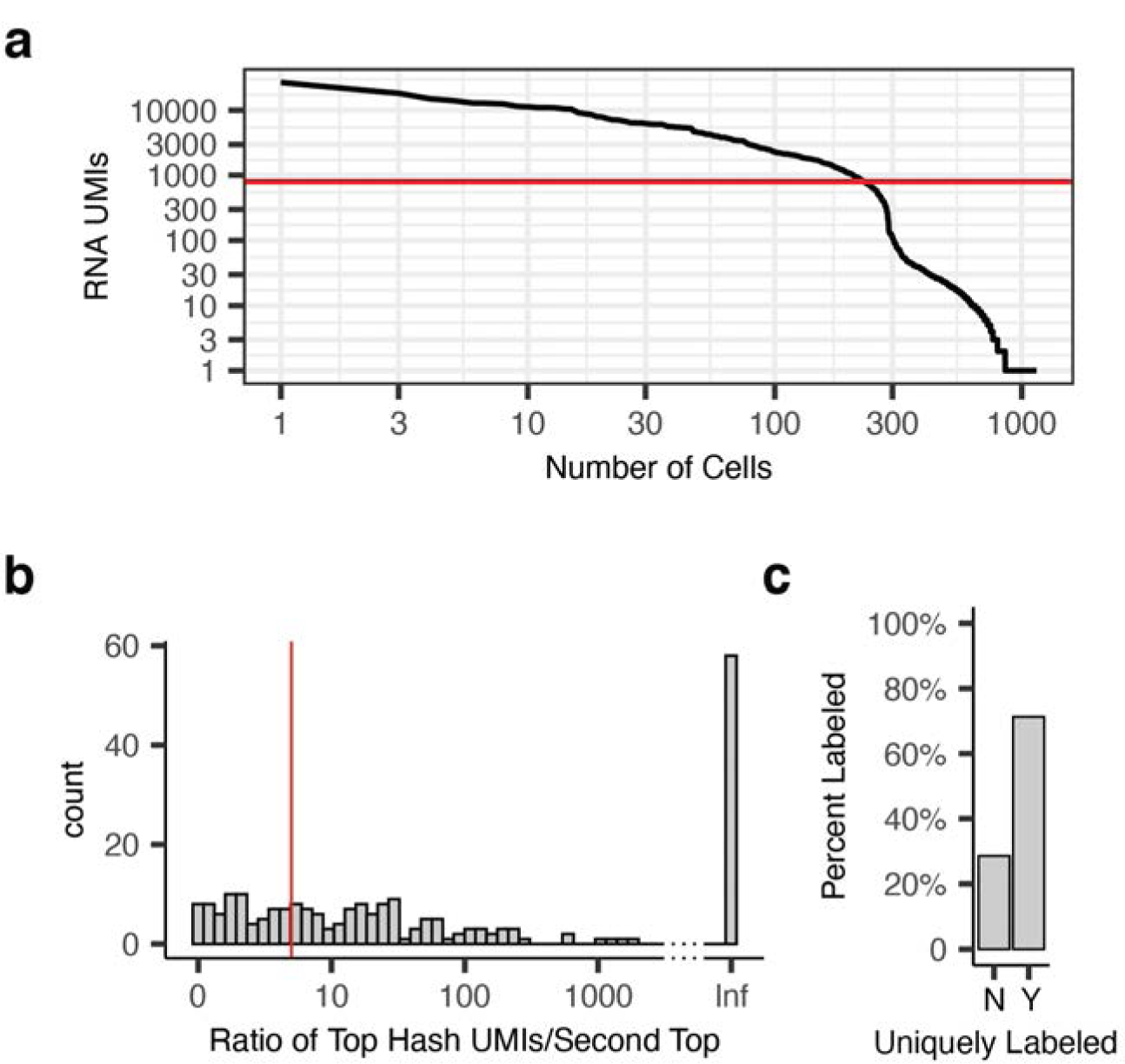
Cells from zebrafish embryos are labeled uniquely with hash oligos. **a**, Ranked plot of the number of unique molecular identifiers per cell (UMIs). Barcodes above the red line (800 UMIs) were called cells. **b**, Enrichment ratio distribution – the ratio between the counts for a cell’s top hash oligo and the second most abundant hash oligo after subtracting background hash molecules. Cells displaying a 5 fold enrichment (red line) of a single hash oligo were deemed uniquely labeled. **c**, Percentage of uniquely labeled cells (Y – uniquely labeled, N – not uniquely labeled).

**Extended Data Figure 2.**
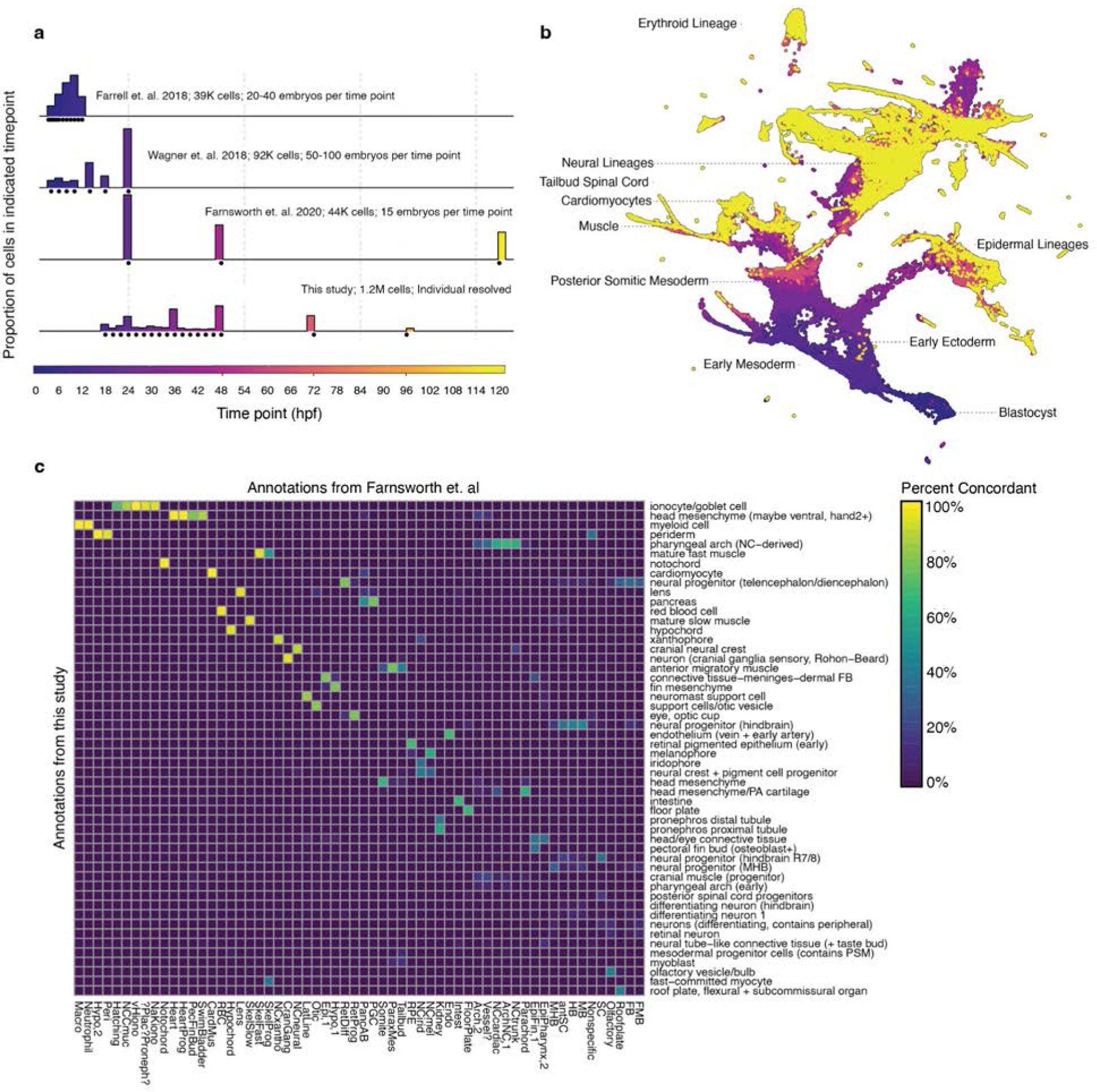
Summary of existing single cell zebrafish atlases and their integration. **a**, Scaled histogram of the proportion of cells deriving from each timepoint in the indicated dataset. Black circles underneath histogram denote distinct collections. Cells and embryos used in each study are indicated as text. **b**, Data integration between Farrell^7^, Wagner^8^ and Farnsworth^17^ datasets spanning 3.3 hours to 24 hours. Cells are colored by collection time matching panel A with select cell lineages labeled. **c**, Heatmap depicting the percentage of each cell type in the Farnsworth^17^ dataset with nearest neighbors in this study at 24 hpf. Columns are annotations from Farnsworth et. al., rows are annotations from this study and each column sums to 100%. Transcriptomes from the two datasets were restricted to a shared set of genes, and downsampled prior to alignment with Seurat.

**Extended Data Figure 3.**
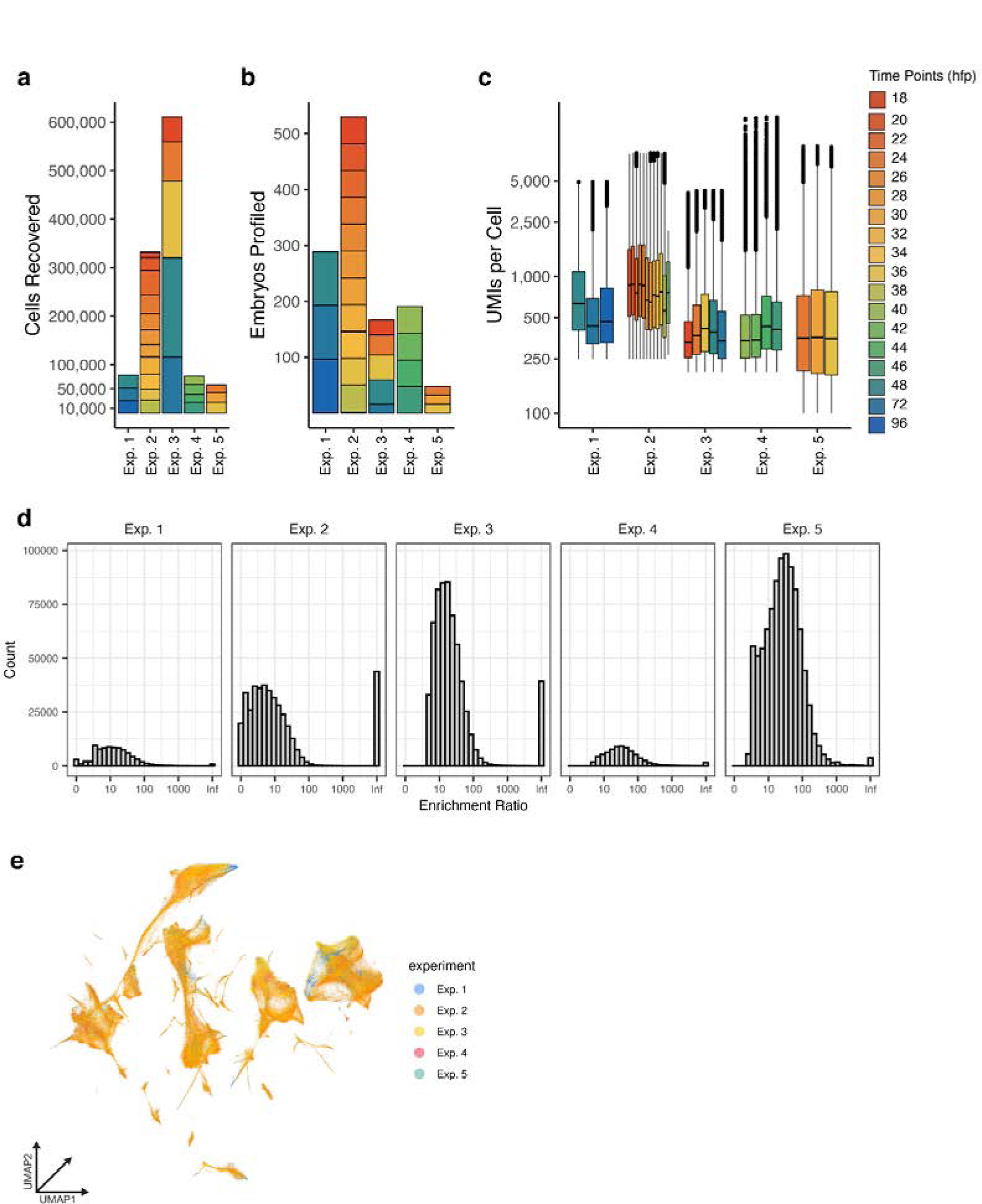
Experimental metrics for reference sci-RNA-seq time series data. **a-c**, Reference dataset summary statistics displaying the number of (**a**) cells, (**b**) embryos and (**c**) UMIs recovered from each experiment. Plots are colored by the time point of embryo collection (panel c) and time points are displayed as hours post fertilization (hpf). **d**, Hash enrichment ratio distribution for cells in the reference dataset. Enrichment ratio was calculated as the ratio of top ranked hash molecules observed in a cell divided by the second most abundant hash molecule after background subtraction. **e**, UMAP embedding in 3-dimensions of the wild-type reference dataset colored by experiment of origin.

**Extended Data Figure 4.**
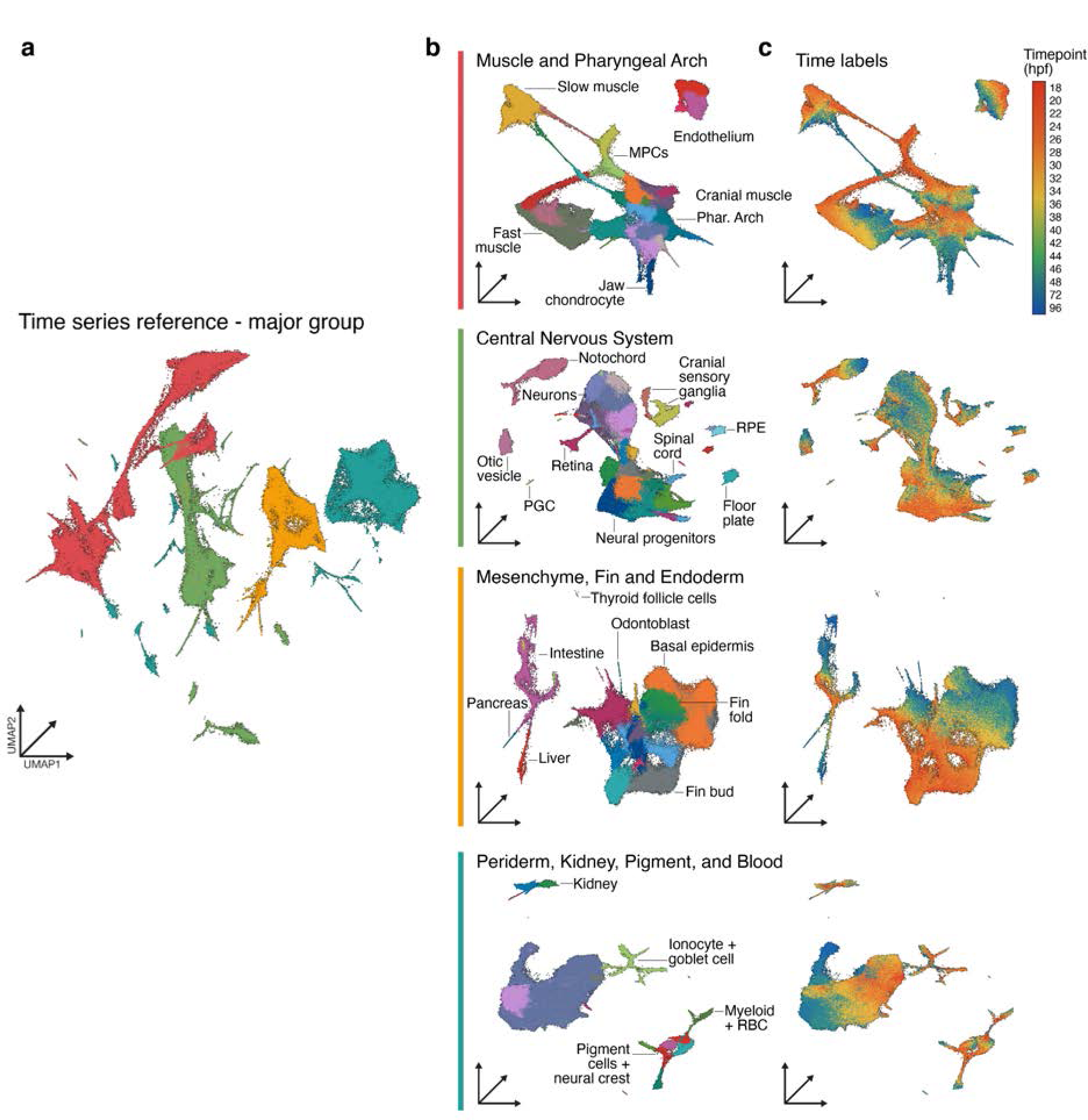
Sub-clustering of major lineages from the reference dataset. **a**, UMAP embedding colored by the 4 major partitions. **b**, Sub-UMAP embeddings of each partition colored by cell type annotation with select cell types labeled. **c**, Sub-UMAP embeddings colored by time point.

**Extended Data Figure 5.**
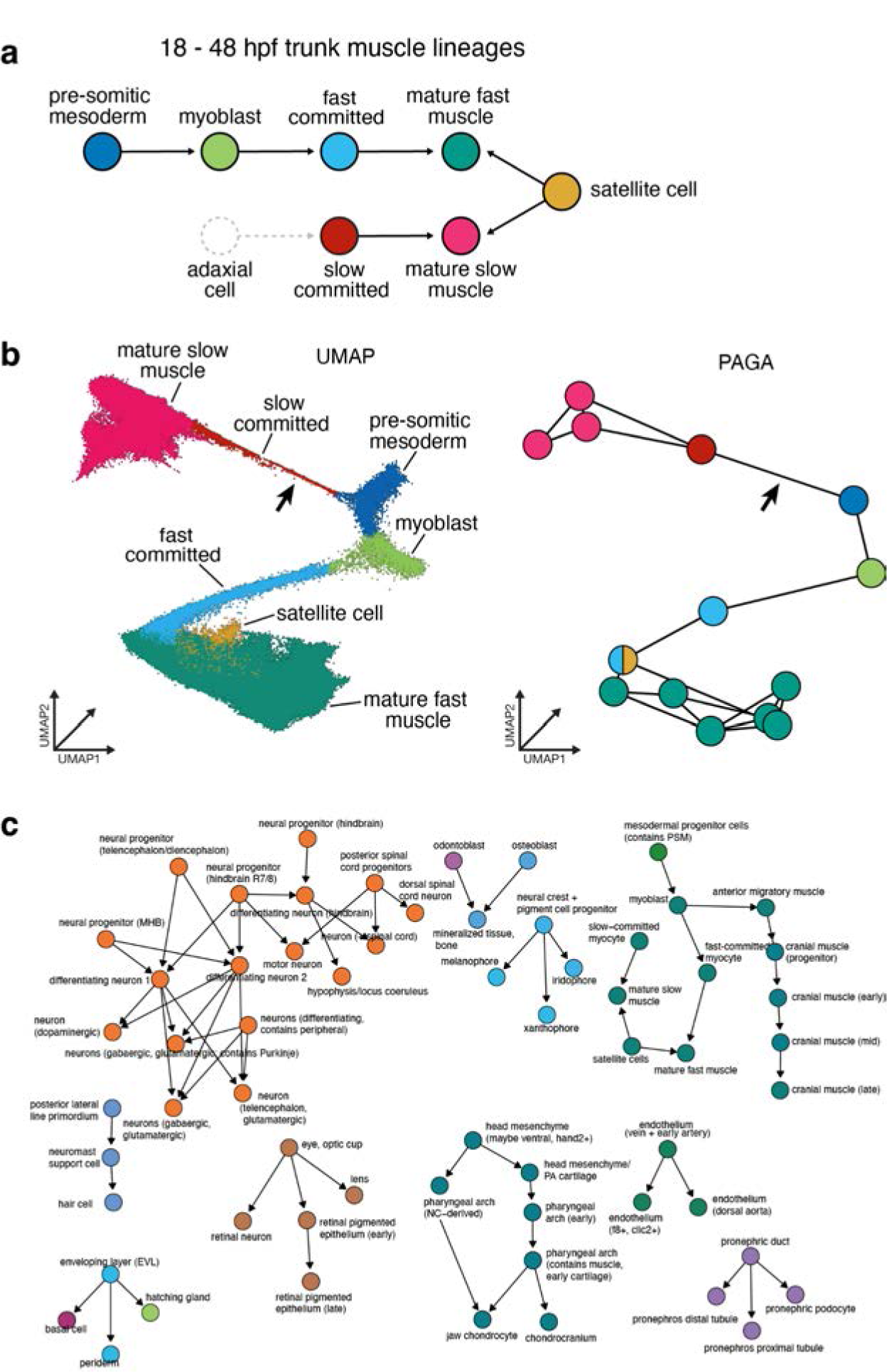
Trunk muscle lineage versus transcriptional relationships. **a**, True lineal relationships between trunk muscle cell types between 18 and 96 hpf. Adaxial cells and linkage to slow committed muscle shown as a dotted line to signify presence prior to 18 hpf (earliest collection). **b**, Transcriptional relationships between cells annotated as trunk muscle types in a UMAP dimensionality reduction plot (3D) and a graph made using the PAGA algorithm ^76^. Arrows indicate connections that exist between transcriptional states. They do not necessarily represent true cell lineage relationships. **c**, A graphical representation of lineal relationships documented in ZFIN between annotated cell types in our reference dataset.

**Extended Data Figure 6.**
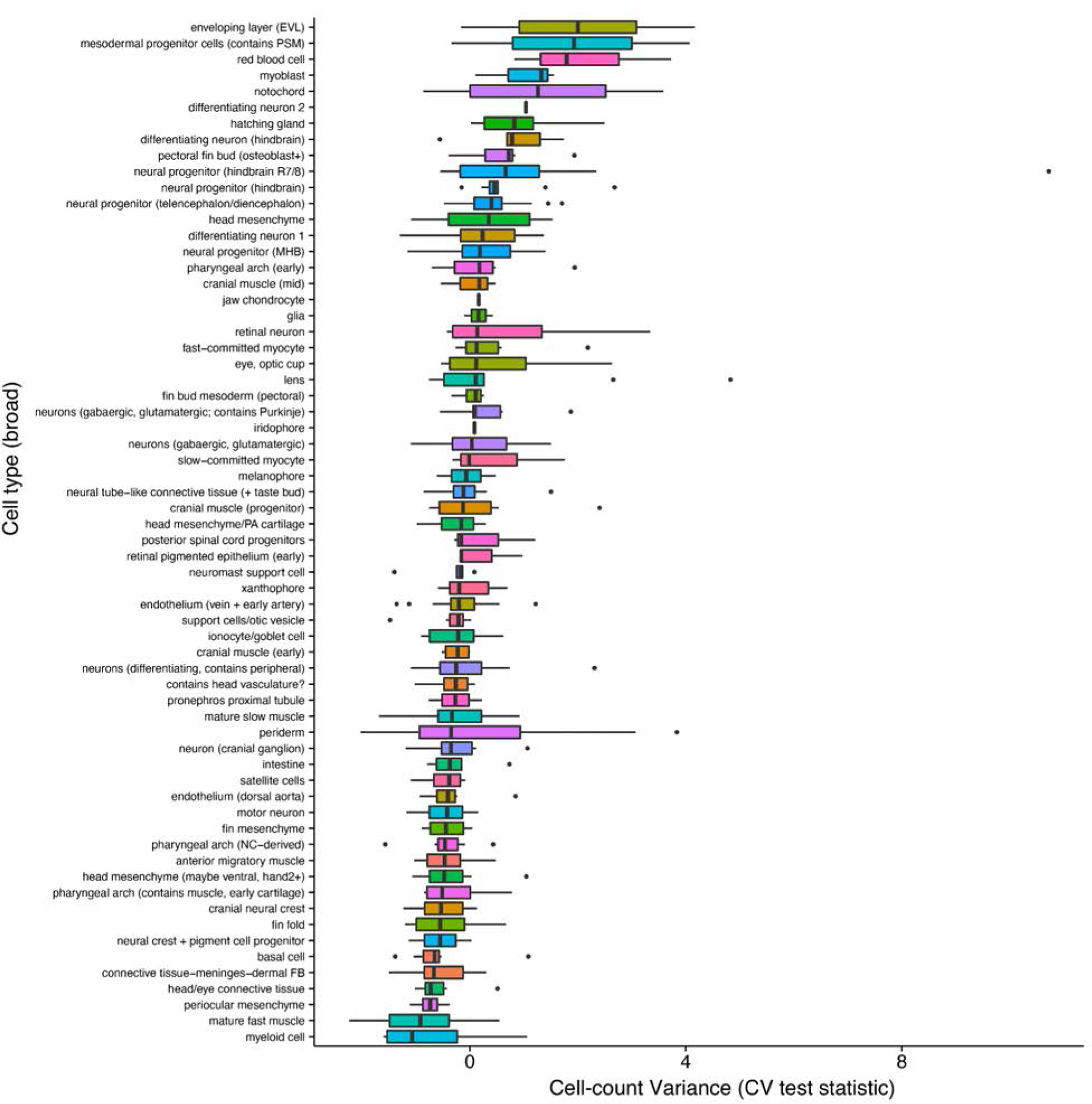
Quantification of variance across all cell types. Cell count mean/variance relationships for all cell types per individual embryo, collapsed by time point and ranked by means. Mean-variance relationships are computed via beta-binomial modeling of the variance, followed by significance testing on the variance observed over the variance expected based on mean cell abundance. Colors denote different cell types.

**Extended Data Figure 7.**
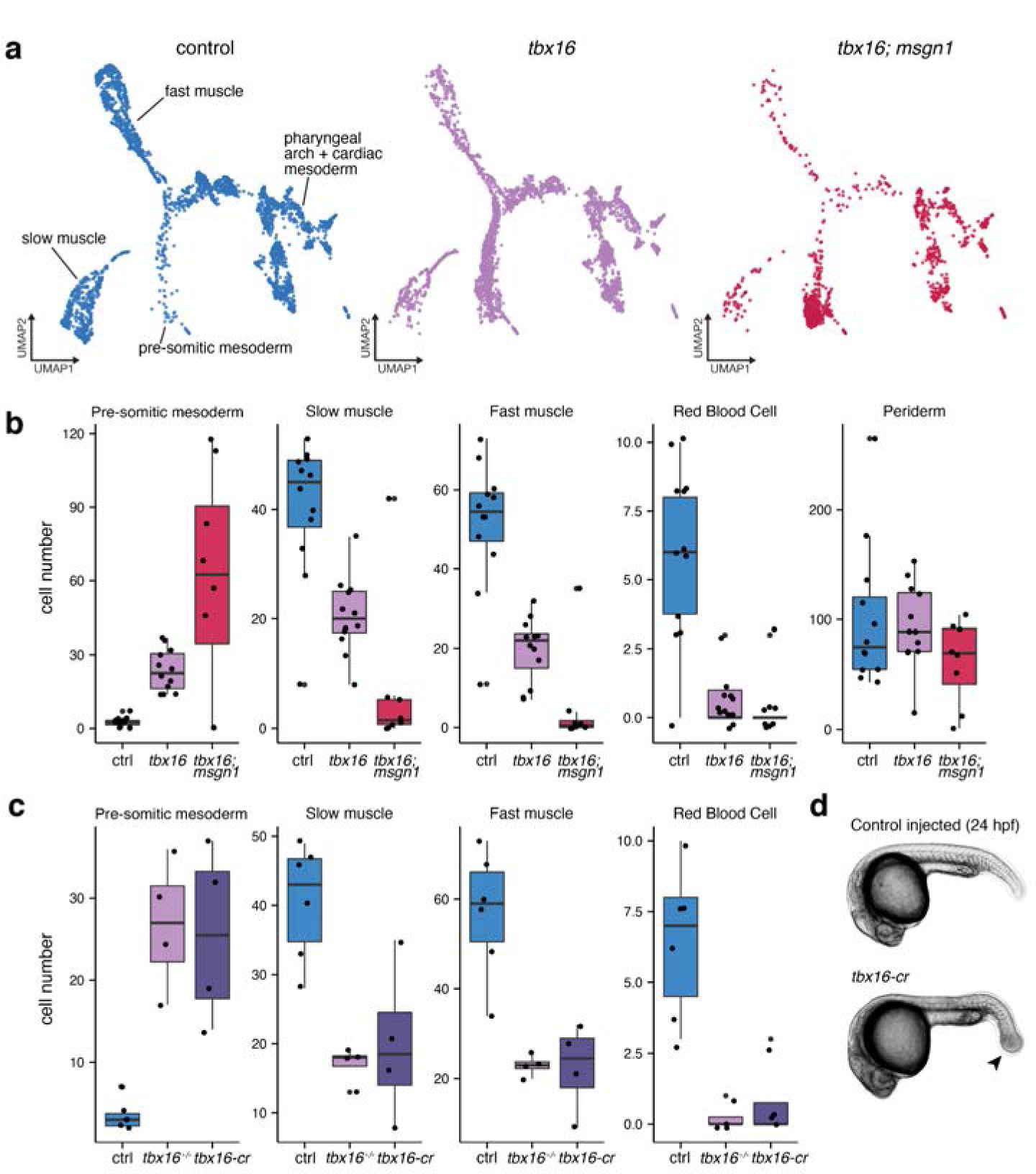
Proof-of-concept whole organism phenotyping with sci-RNA-seq for well-characterized mutants and crispants. **a**, UMAP embedding of the mesodermal trajectory from whole organism sci-RNA-seq (n = 6,443 cells; total n = 29,800 cells). Plots are faceted and colored by their perturbation (control, *tbx16* (mutants and crispants), and *tbx16*;*msgn1* crispants). **b**, Boxplot of the size factor-normalized counts of each cell type recovered from individual embryos split out by perturbation. Size factor-normalized counts from each embryo are displayed as points over each boxplot. Cell types displayed are those predicted to have differential abundances in response to *tbx16* or *tbx16;msgn1* loss of function except periderm, which is not changed. **c**, Cell count boxplots for control embryos relative to *tbx16*-/- or *tbx16* crispants (*tbx16-cr)*. **d**, Representative images of control injected and *tbx16-cr* embryos at ∼24 hpf. Characteristic “spadetail” indicated by arrow.

**Extended Data Figure 8.**
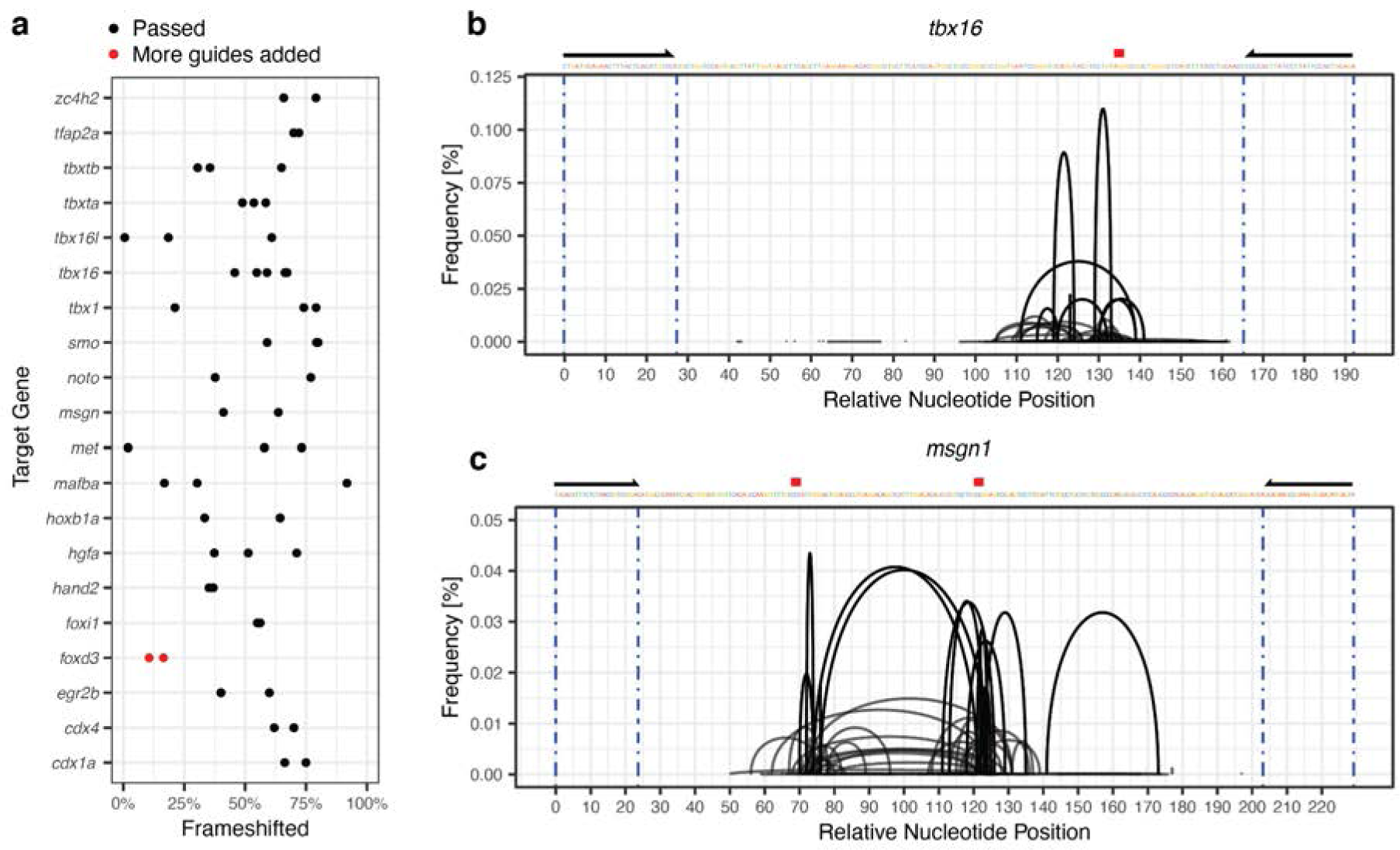
Ascertaining gRNA mutagenesis rate using multiplex PCR. **a**, Percentage of frame shifted amplicons amplified from CRISPR-Cas9 edited zebrafish. Extra guides were added for Foxd3 (red points) due to a low editing rate, and the absence of the expected phenotype. **b-c**, Frequency of the cut sites detected within amplicons for Tbx16 (**b**) and Msgn1 (**c**). Black primers flanking the targeted region denote primers used for amplification of the amplicon. Protospacer adjacent motif displayed as a red box above the sequence. Mapping, analyses and plots deployed the ampliCan software package in R ^77^.

**Extended Data Figure 9.**
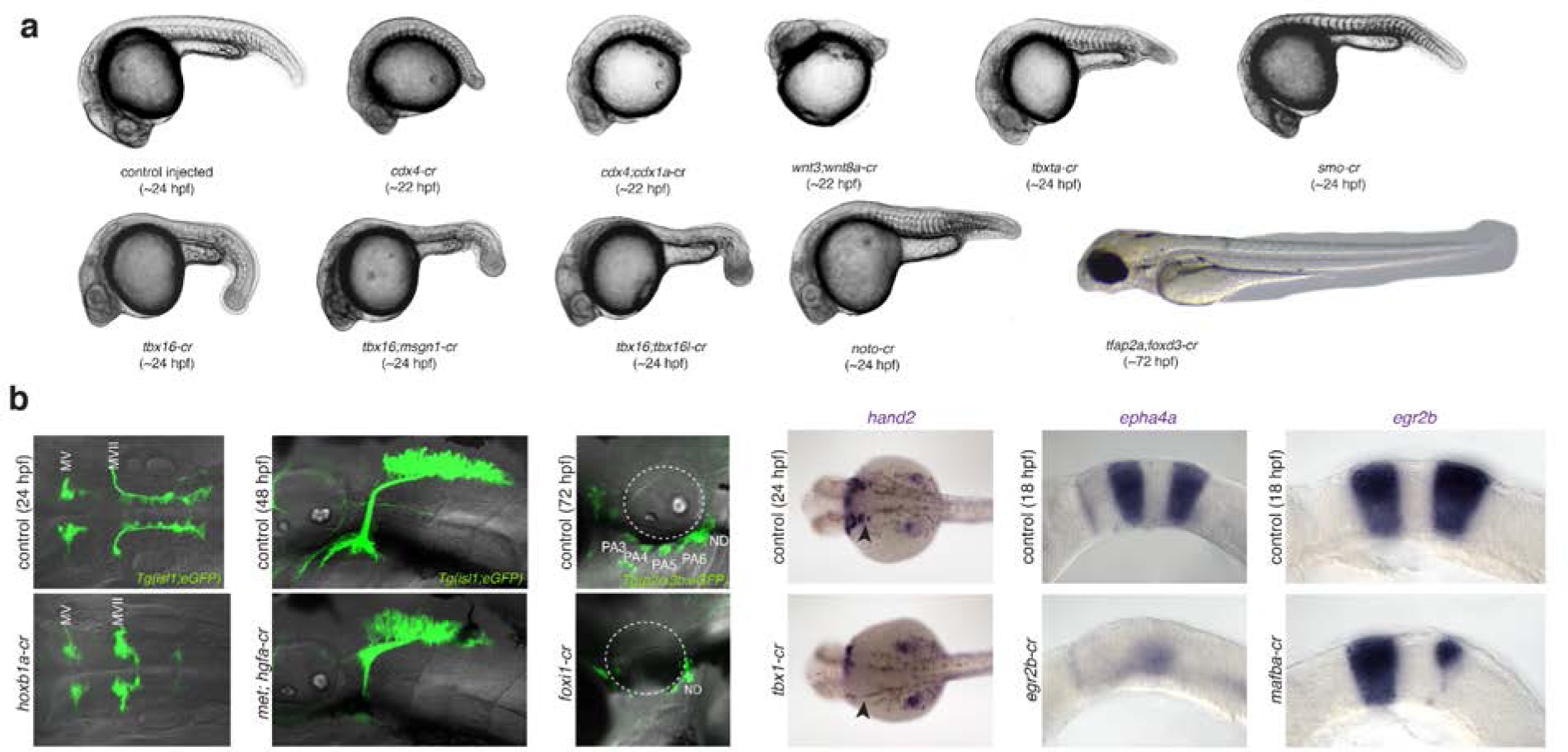
Phenotypic confirmation of F0 CRISPR-Cas9 injections. **a**, In addition to mutagenesis efficiency, gRNA sets were selected for their ability to generate phenotypes in F0 animals that resembled published null phenotypes. Representative images are labeled by their approximate developmental time and perturbation. **b**, For embryos where phenotypes were not apparent via whole mount, brightfield views, we evaluated the perturbation using appropriate transgenic lines or ISH. ISH target genes, perturbations, approximate time points, and anatomical landmarks are labeled (MV, trigeminal motor neurons; MVII, facial motor neurons; white dotted circle, ear; black arrow, posterior pharyngeal arches; PA#, pharyngeal arch number; ND, nodose ganglion).

**Extended Data Figure 10.**
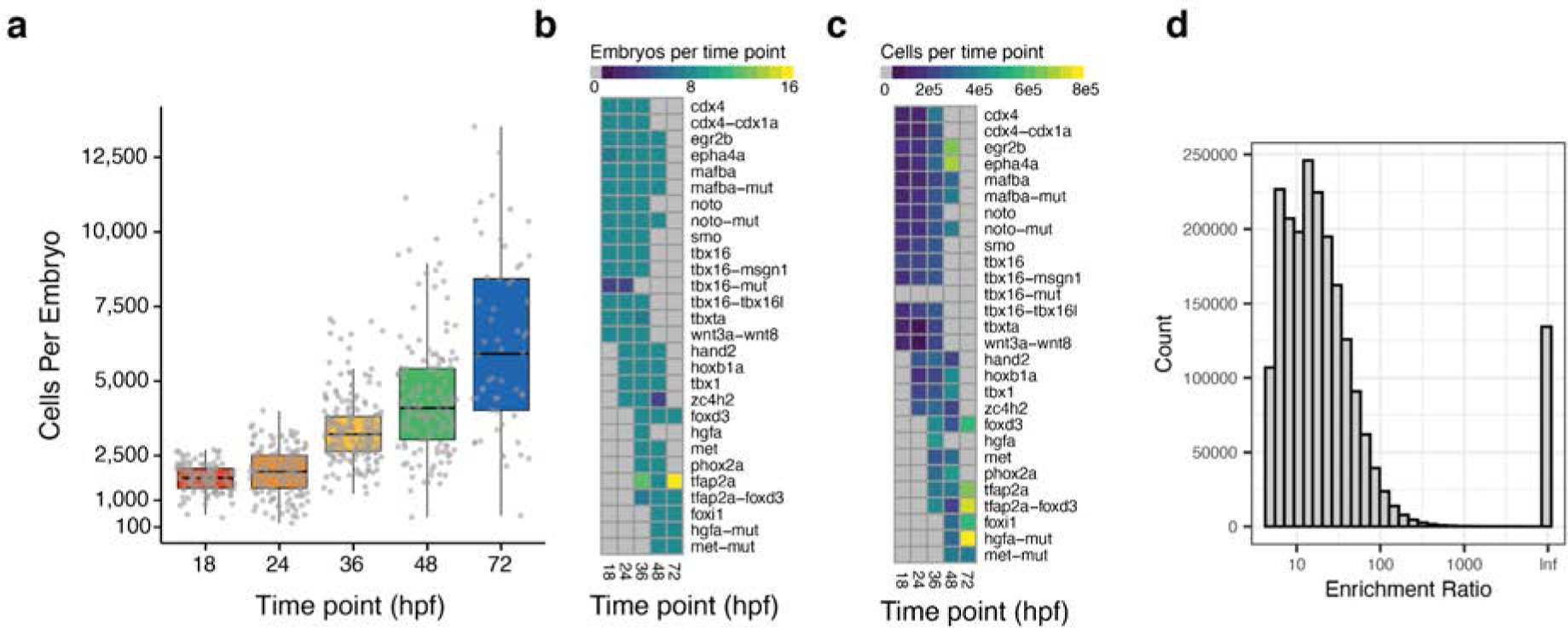
Quality control metrics for crispant scRNA-seq data. **a**, Number of cells recovered per mutant embryo. Each gray point is an individual embryo that is summarized by the boxplot. Prior estimates suggest that a 24 hpf embryo has ∼25,000 cells ^78^; based on this, we estimate a 5-10% recovery per embryo. **b**, Heatmap displaying the number of embryos collected per perturbationxtime point combination. **c**, Heatmap of the number of cells collected from each perturbation
xtime point combination. **d**, Hash enrichment ratios for cells in the mutant dataset. Enrichment ratio was calculated as the ratio of top ranked hash molecules observed in a cell divided by the second most abundant hash molecule after background subtraction.

**Extended Data Figure 11.**
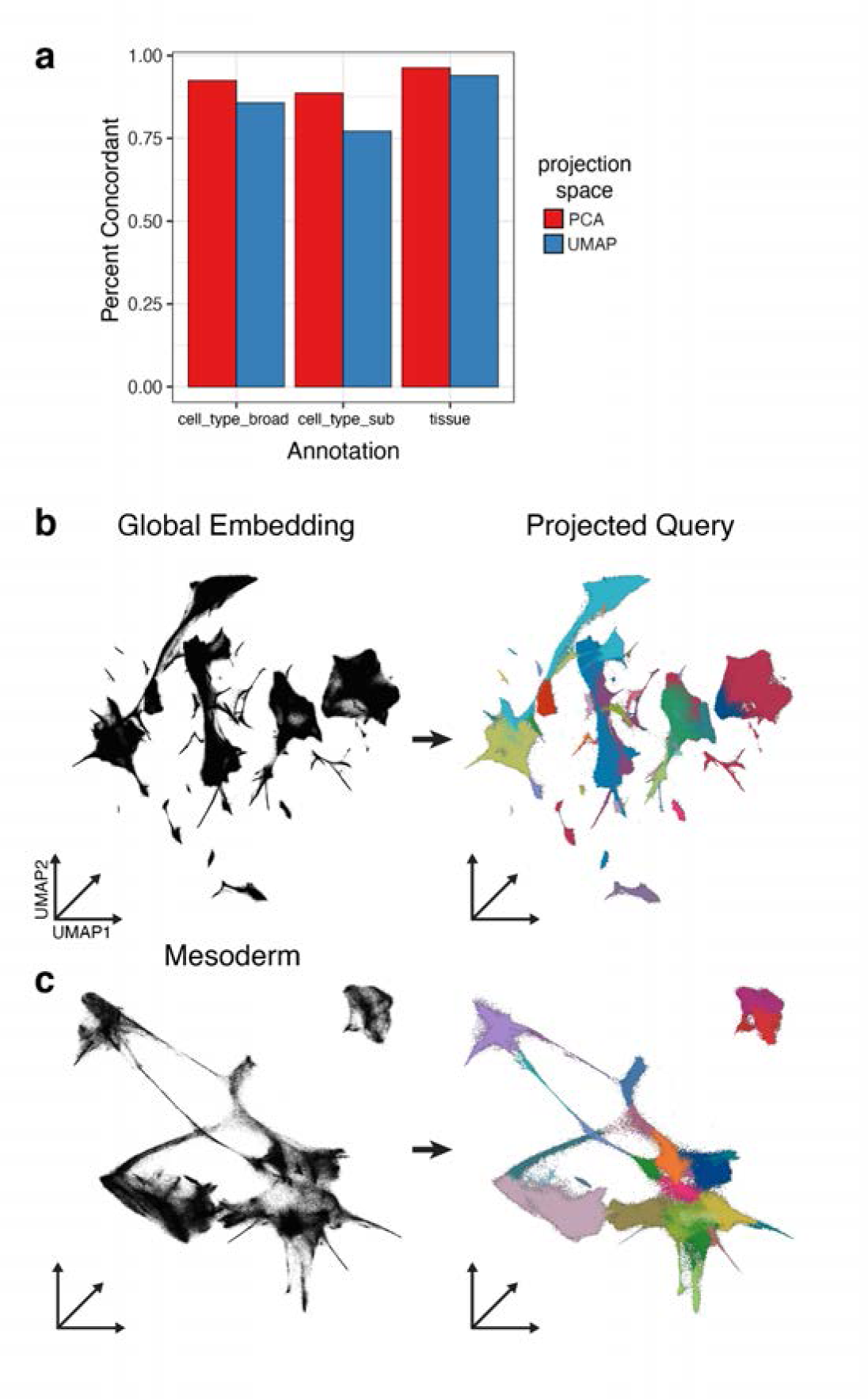
Query to reference dataset projection. **a**, Manually annotated wild-type reference cells were split 80:20. The 80 percent split was used as input for PCA, followed by UMAP. The 20 percent split was projected using the same transformations and labels were transferred in PCA space or UMAP space. Annotation labels were then transferred in either PCA space (red) or UMAP space (blue). Labels were deemed concordant if manual annotation matched the projected transfer annotation. **b-c**, Schematic depicting label transfer of the tissue annotation (**b**), followed by transfer of broad cell type annotation in the mesoderm tissue (**c**).

**Extended Data Figure 12.**
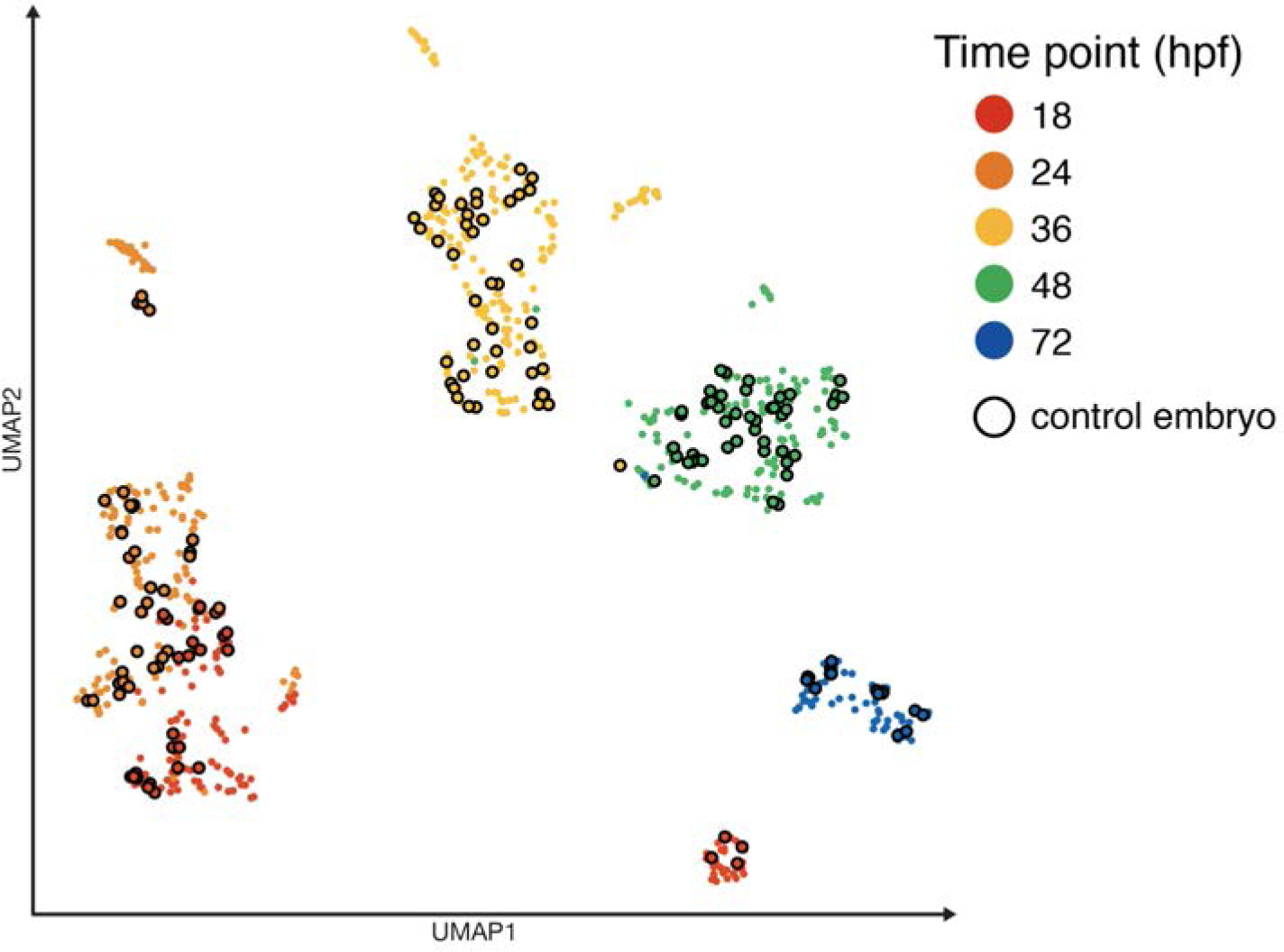
A low dimensional representation of embryo composition for control and perturbed embryos. A UMAP plot where each point represents the cell type abundance composition (i.e. counts) for a single embryo, colored by collected time point. Perturbed embryos lack borders, and points with a black border is a control injected or null wild type sibling embryo.

**Extended Data Figure 13.**
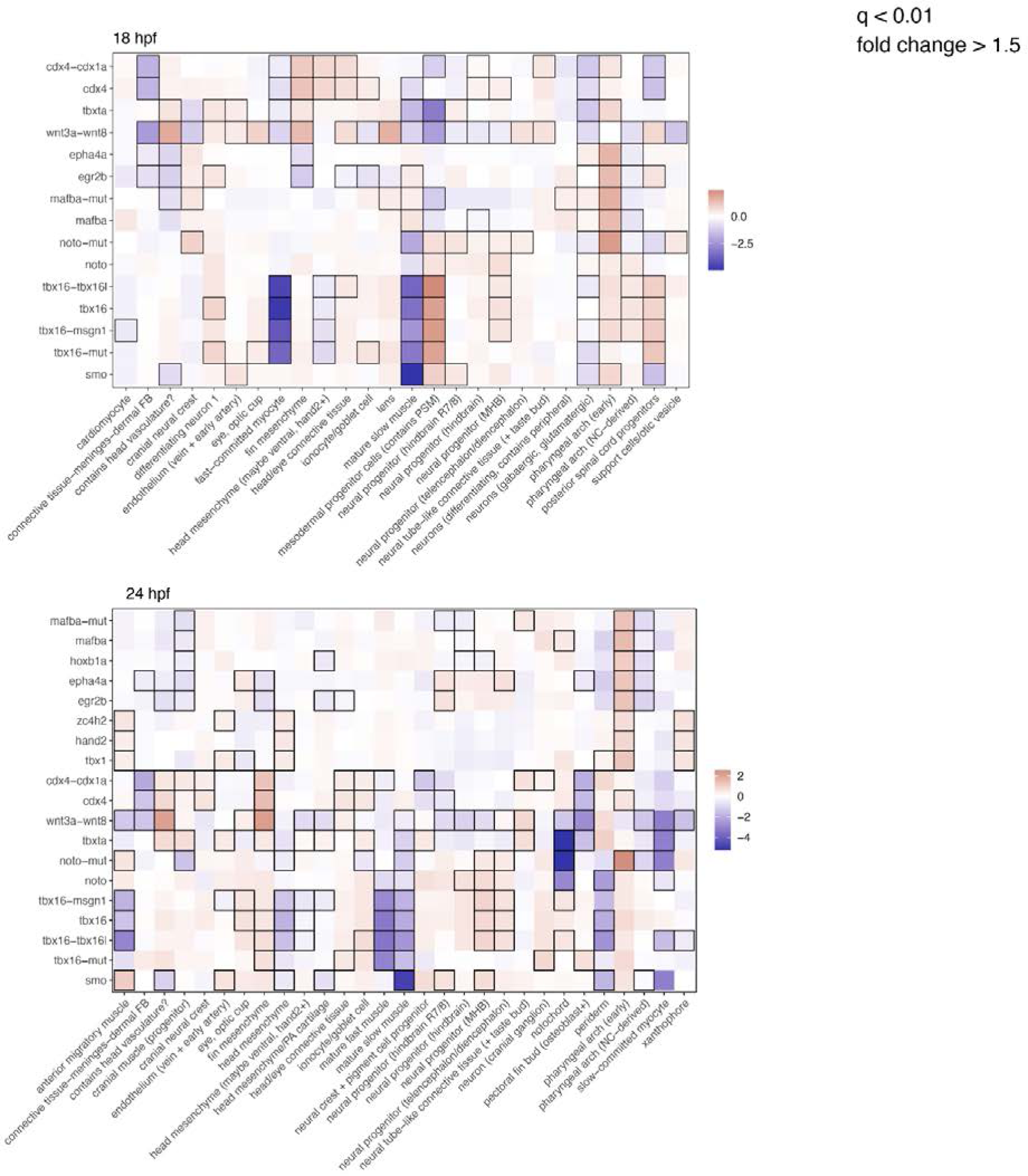
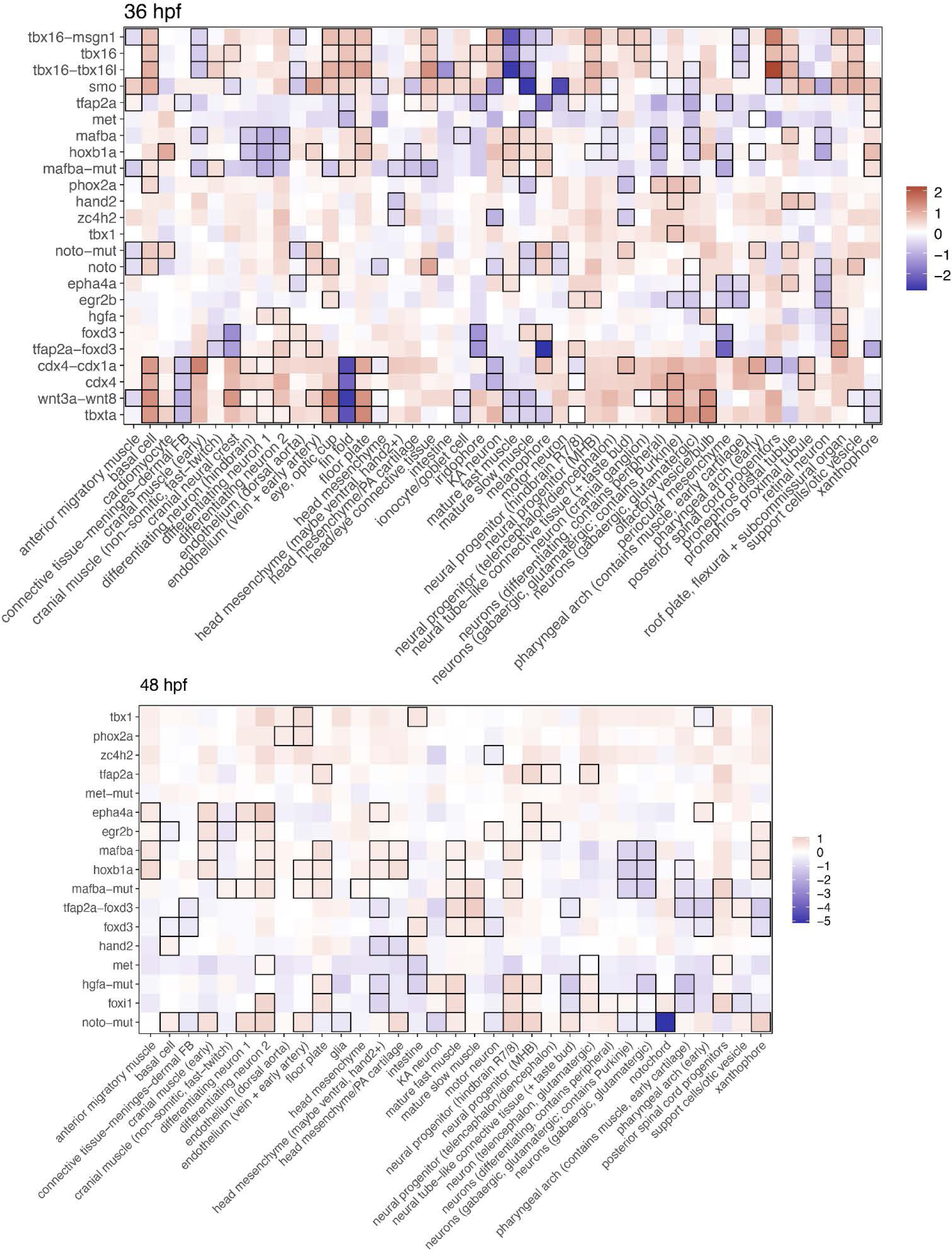
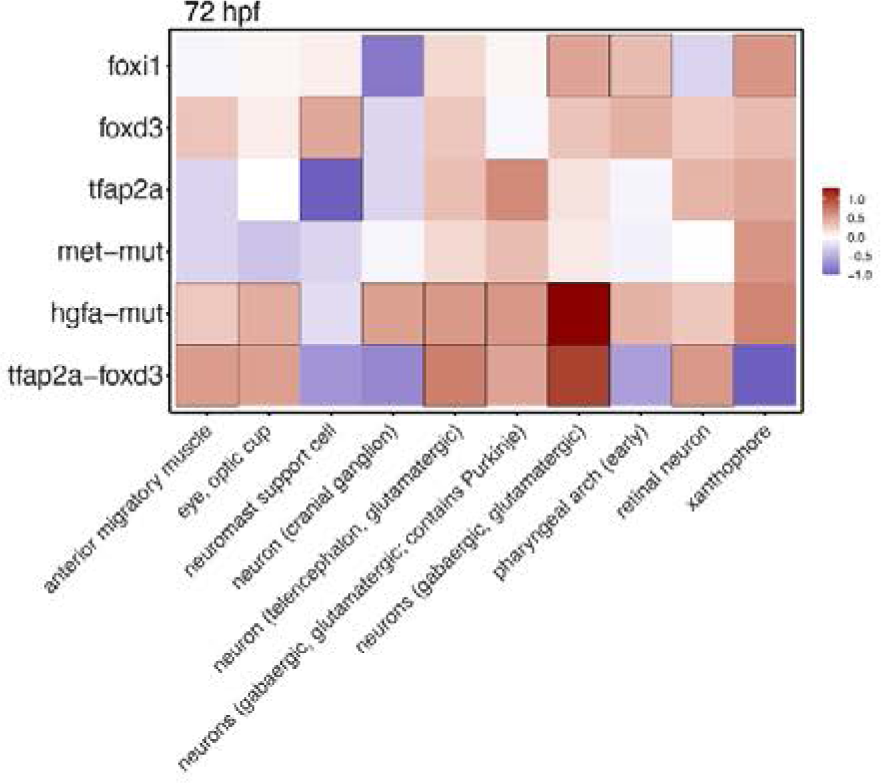
Differentially abundant cell types across all perturbations and timepoints. Heatmaps representing the log_2_ fold-change in abundance of each perturbation relative to control-injected or wild type cells, for each cell type. Boxes indicate significant changes as determined using a beta-binomial regression modeling approach (*q-value* < 0.01, fold change > 1.5).

**Extended Data Figure 14.**
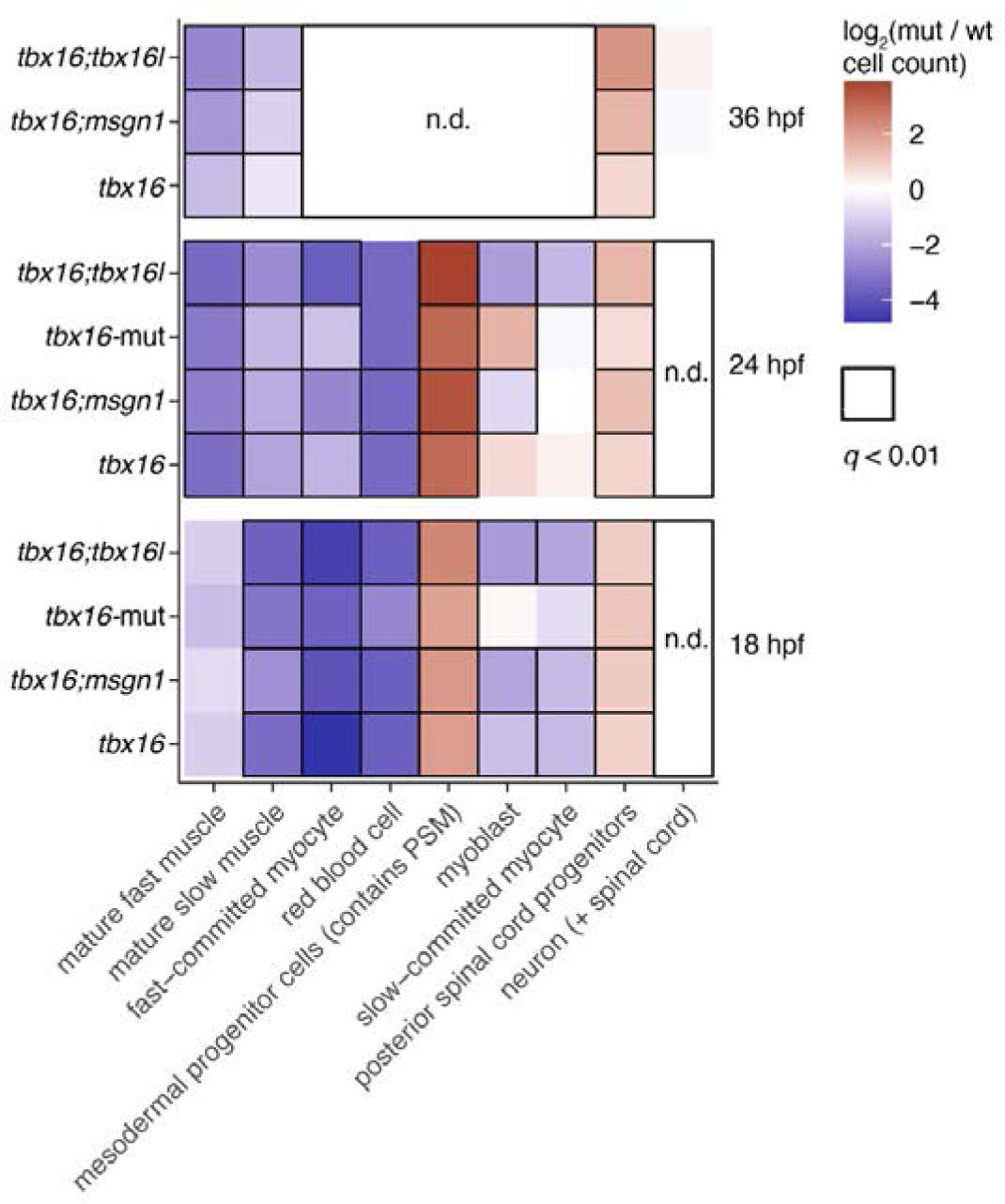
Differential cell type abundance across t-box groups in the mesoderm and spinal cord neurons. A subset of trunk muscle and spinal cord neuron cell types for each of four perturbations relative to control embryos at matched timepoints: *tbx16, tbx16; msgn1, tbx16; tbx16l,* and *tbx16^-/-^*. Black boxes indicate significance (*q* < 0.01, beta binomial regression; n.d. - no cells of this type detected at these stages).

**Extended Data Figure 15.**
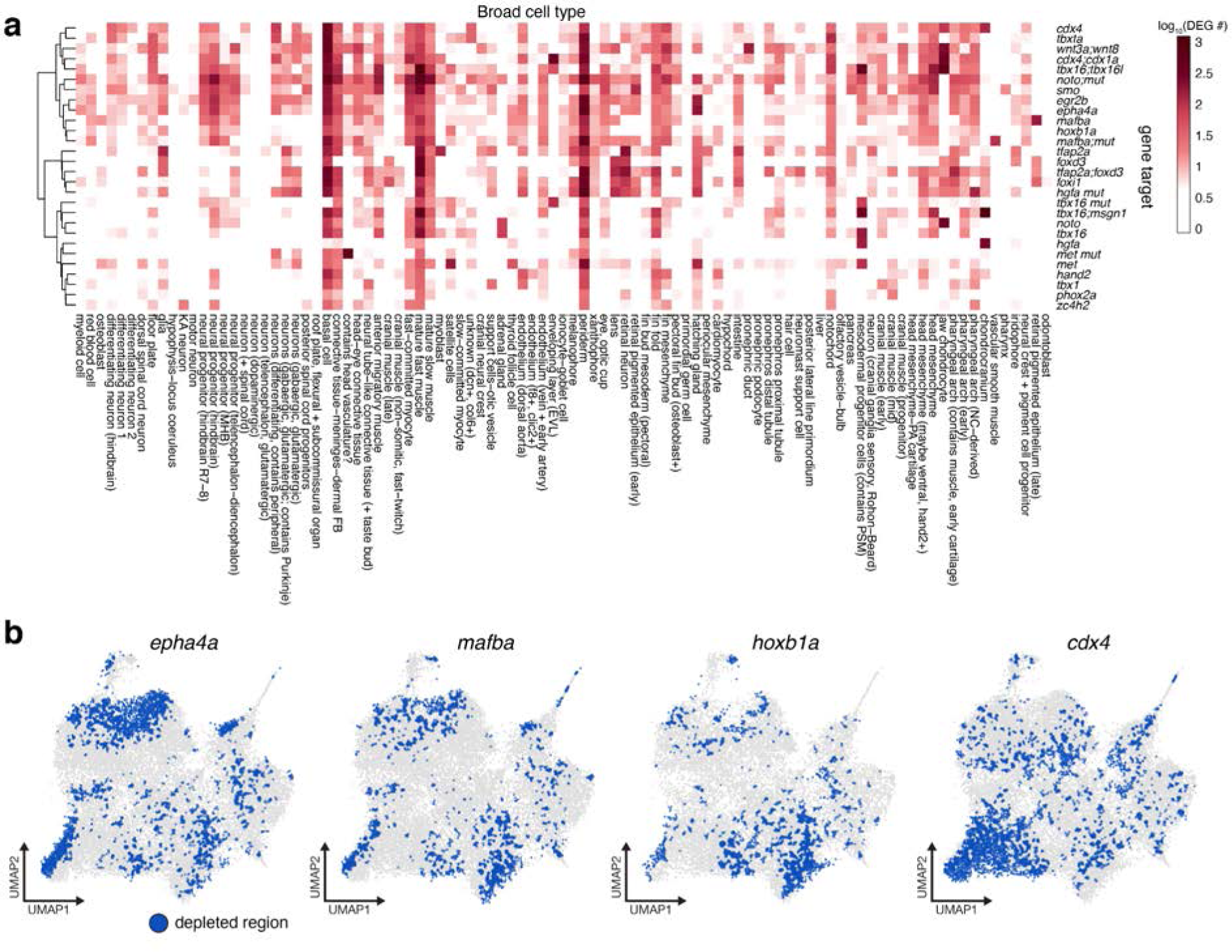
Transcriptional responses to genetic perturbations across targets and cell types. **a**, A heatmap displaying the number of differentially expressed genes (*q* < 0.05) for each broad cell type, across all perturbations. Numbers are displayed in log_10_(x +1). **b**, UMAP plots in which all neural progenitor cells are grey, and blue cells are control cells that are determined with the Getis-Ord test to have neighbors depleted for the perturbed cell type, termed “cold spots” for a selected set of perturbations known to affect hindbrain development.

**Extended Data Figure 16.**
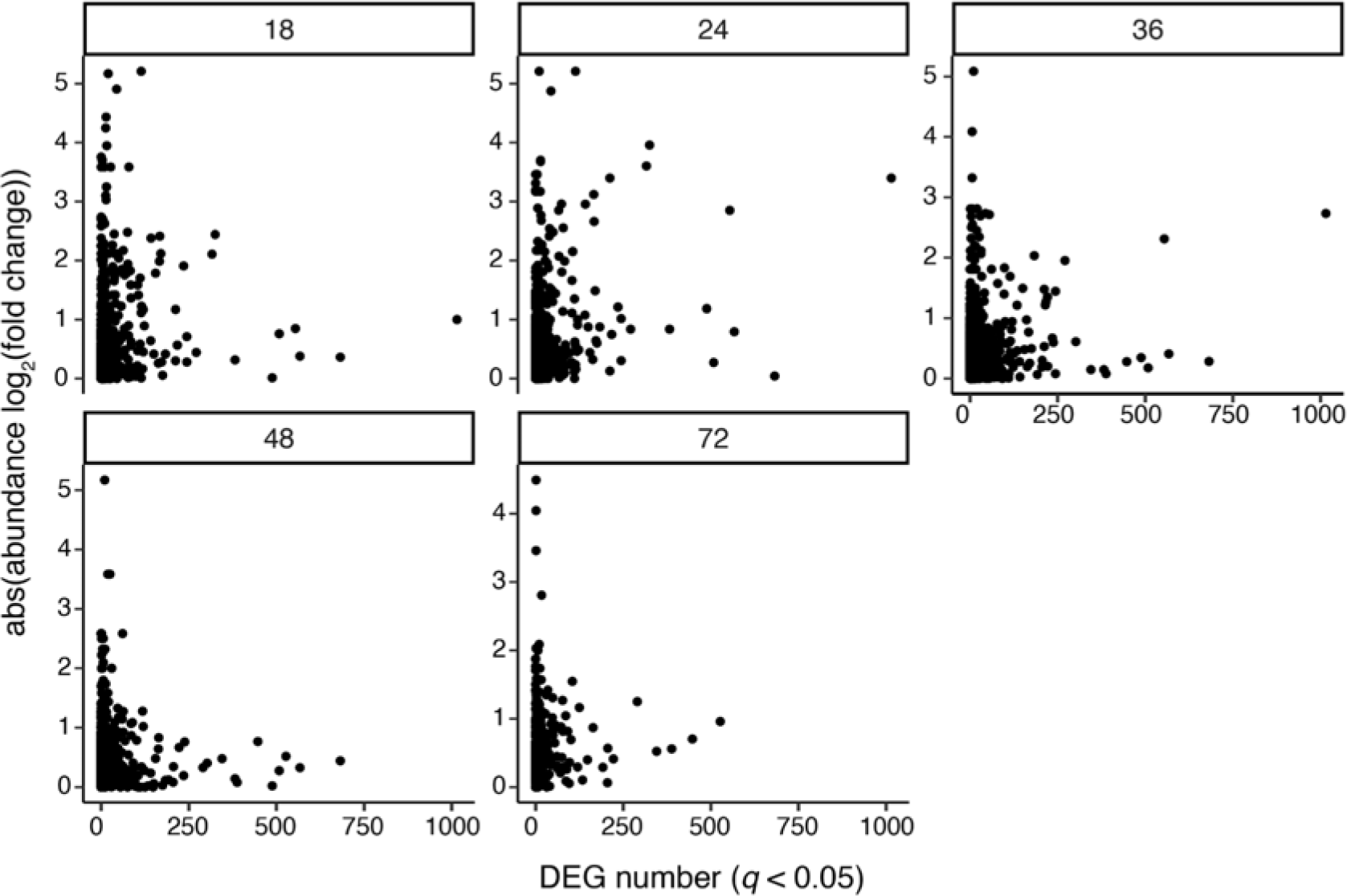
Comparison of perturbation induced DEGs to cell type abundance changes. Scatter plots displaying the number of significant, differentially expressed genes between perturbed cells and control cells (y-axis), versus the absolute fold change in cell type abundance between perturbed and control (x-axis). Each point represents a unique cell type, perturbation pair, and plots are faceted by time point. Cell type specific, differentially expressed genes resultant to perturbation pairs are not associated with changes in cell type abundance.

**Extended Data Figure 17.**
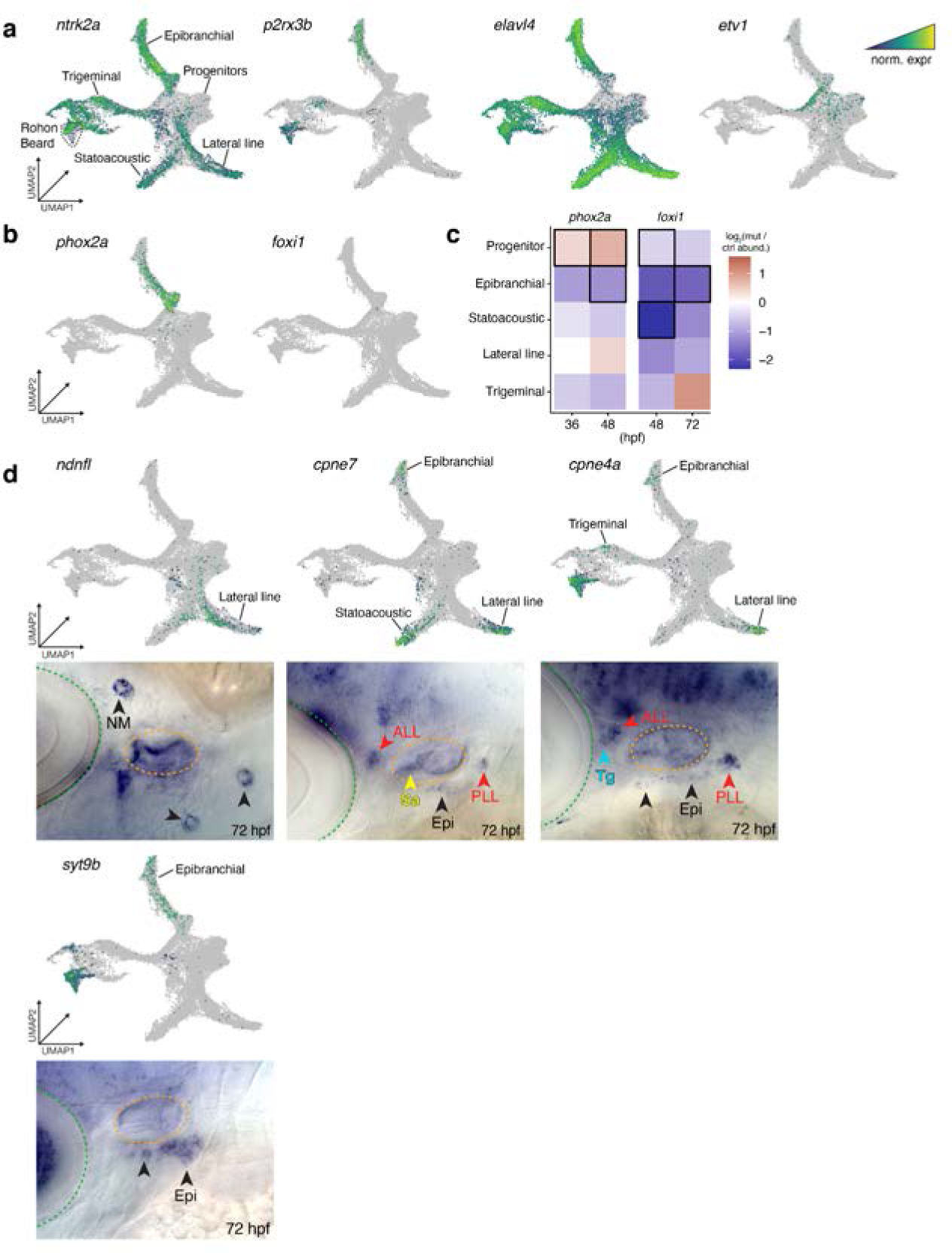
Subtype-specific gene expression in the cranial sensory ganglia. **a**, A UMAP plot where each cell is colored by its mean, normalized expression of neuronal or cranial ganglia markers: *ntrk2a, p2rx3b, elavl4, etv1.* **b**, A UMAP plot where each cell is colored by its mean, normalized expression of *phox2a* or *foxi1.* **c**, Cranial ganglia sensory neurons and their cell abundances (log2, size factor normalized) relative to control embryos at two timepoints for *phox2a* and *foxi1* crispants. Black squares indicate significance (*q* < 0.01). **d**, UMAP expression plots and corresponding ISH with a lateral head view and cranial ganglia labeled. Eye is marked by a green dotted line, ear is marked by an orange dotted line. (LL, lateral line ganglia; PLL, posterior lateral line ganglion; ALL, anterior lateral line ganglion; Tg, trigeminal ganglion; Epi, epibranchial ganglia; Sa; statoacoustic ganglion).

**Extended Data Figure 18.**
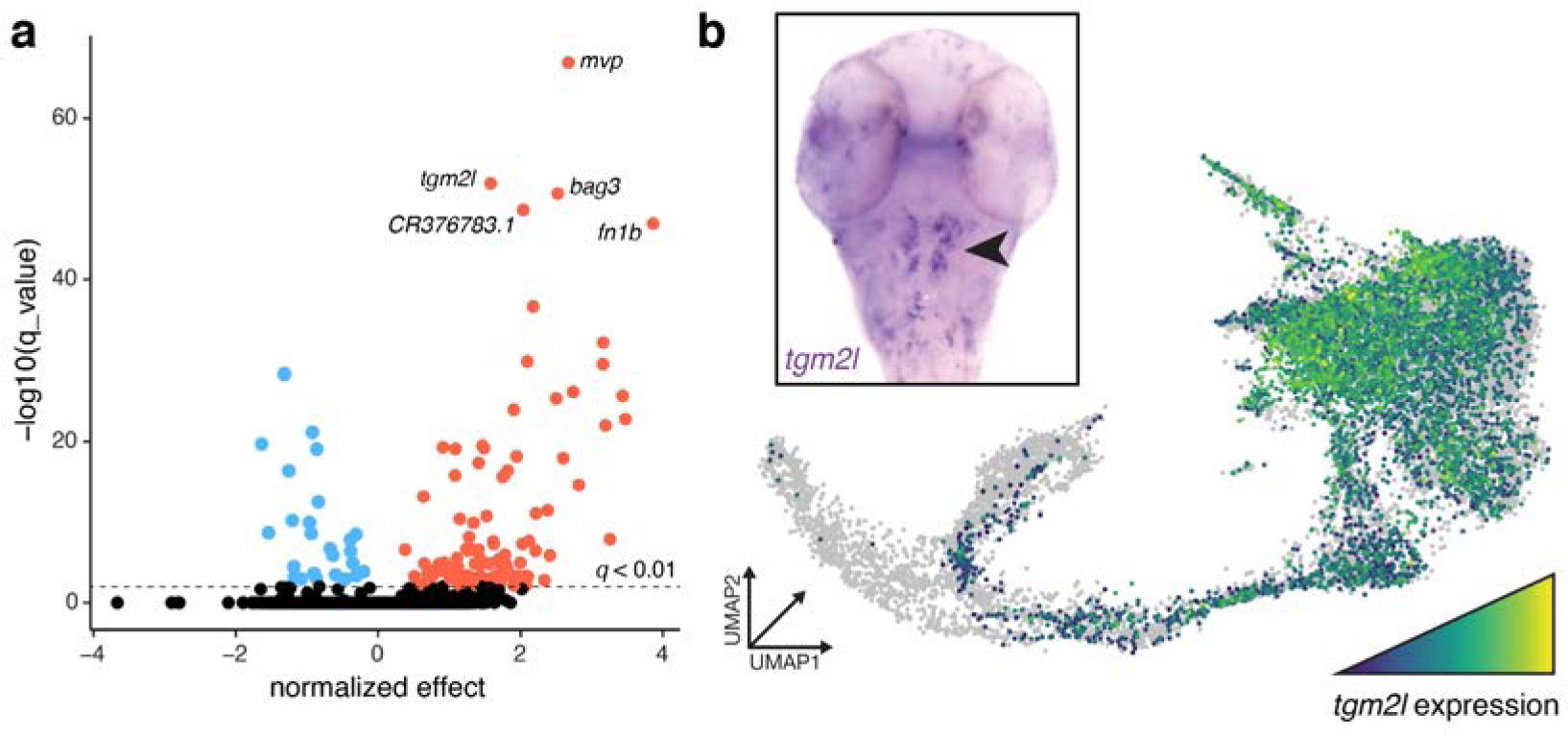
Notochord and PC transcriptome comparison. **a**, A volcano plot representing the differentially expressed genes between *tbxta* PC cells and control notochord sheath and PC cells at 36 hpf. Genes that are enriched in *tbxta* cells are red and genes enriched in control cells are blue. Genes with a *q-*value < 0.01 are black. The top 5 differentially expressed genes are labeled. **b**, A UMAP plot colored by the expression of *tgm2l* in the notochord of control and *tbxta*. *tgm2l* is enriched both in *tbxta* cells (*q* = 4.5E-61) relative to controls at 36 hpf and in the region of the UMAP predicted to be enriched for parachordal cartilage cells. Expression of *tgm2l* via *in-situ* hybridization in PC in a wildtype embryo at 36 hpf (black arrowhead); *tgm2l* is not detectable in notochord cells.

**Extended Data Figure 19.**
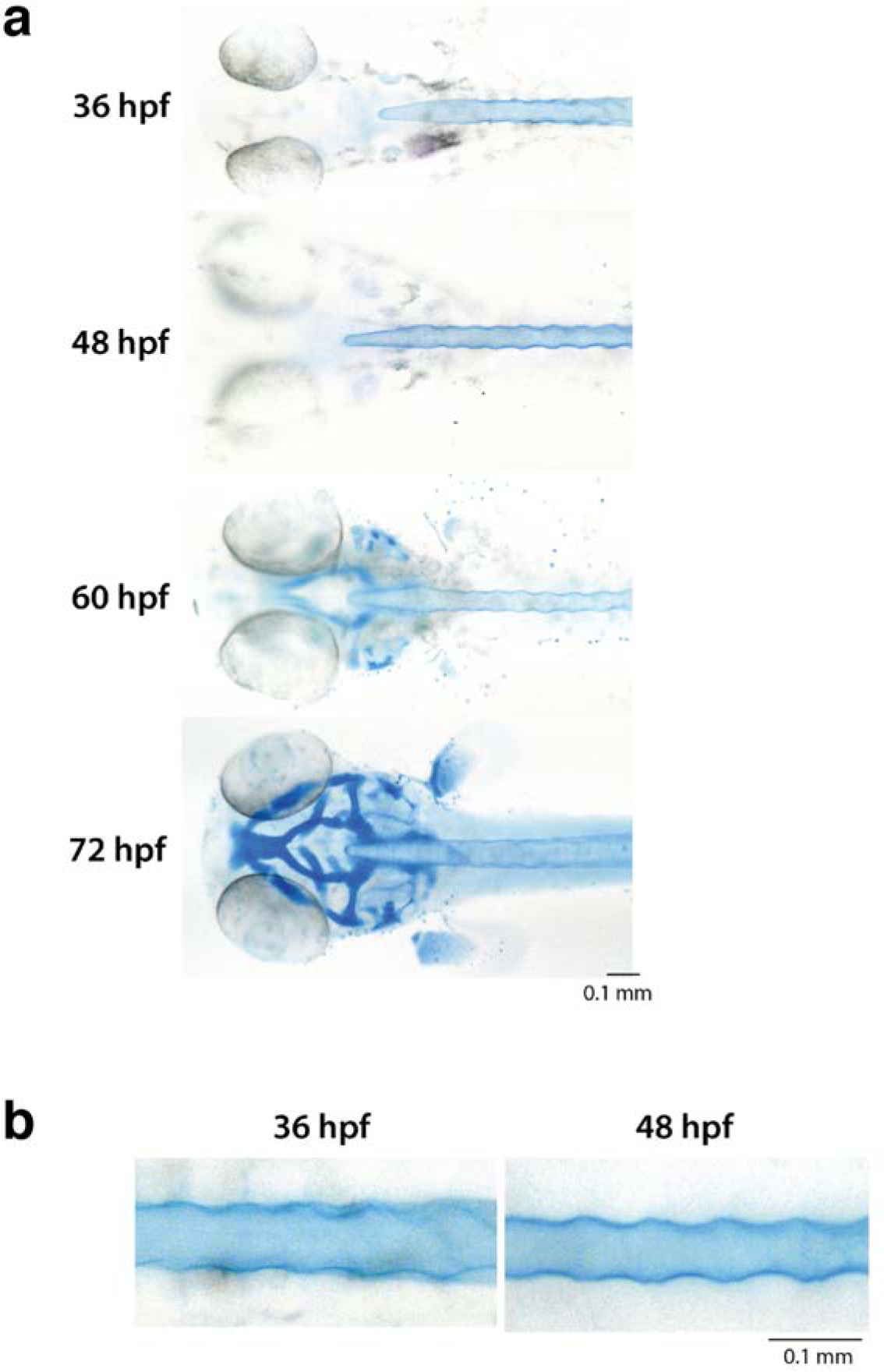
Anterior cartilage development during late embryonic stages. **a,** Anterior dorsal views of alcian blue-stained zebrafish embryos from 36 hpf to 72 hpf. **b**, Dorsal images of alcian blue-stained notochords at 36 and 48 hpf.

**Extended Data Figure 20.**
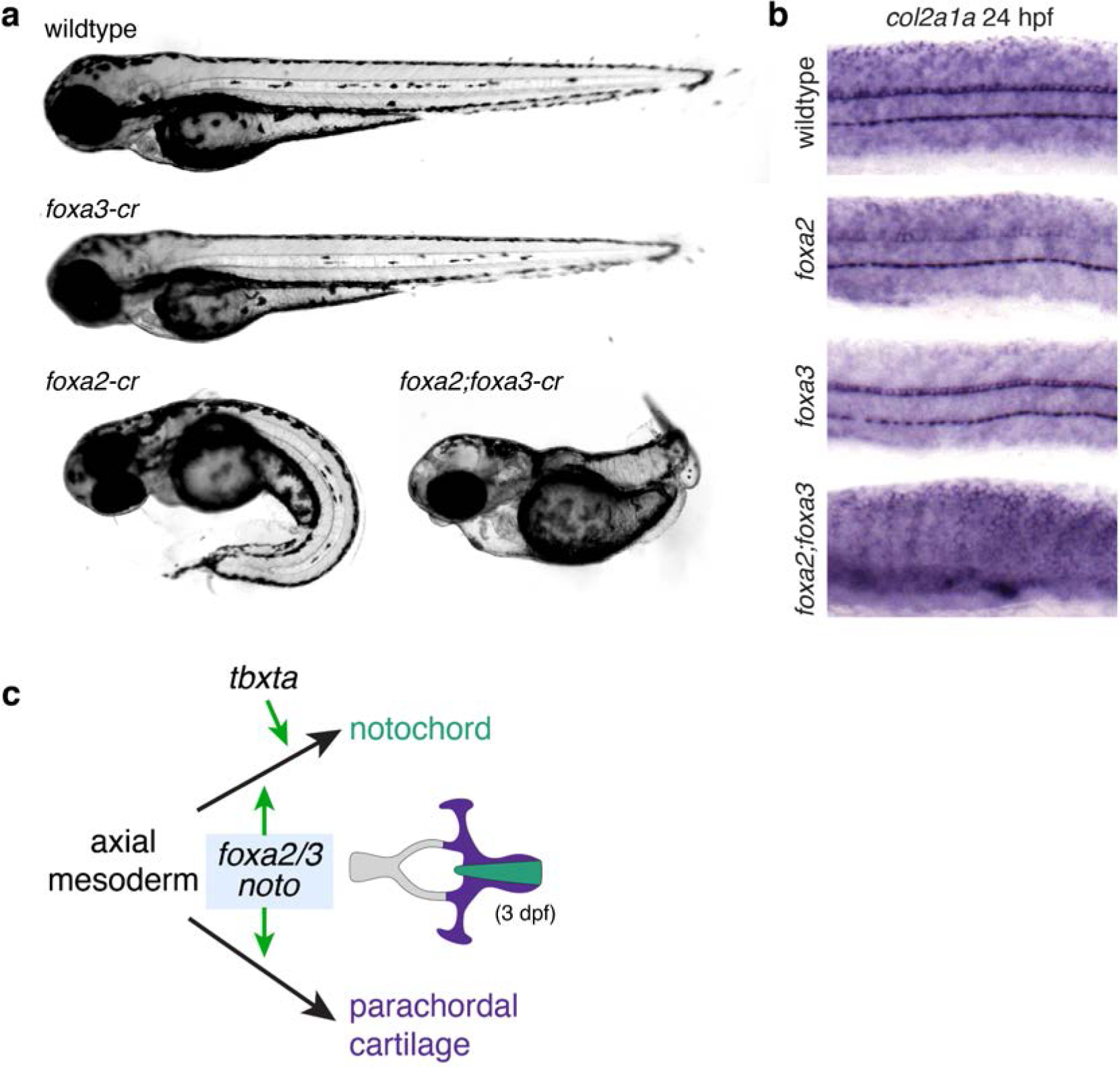
Roles for *foxa3 and foxa3* during notochord development. **a**, Representative images of *foxa2*, *foxa3*, or *foxa2;foxa3* crispants at ∼48 hpf. **b**, Trunk sections for *col2a1a in situ* hybridizations. No notochord cells are present in double *foxa2*;*foxa3* crispants. **c**, An updated model of the independent genetic requirements for PC and notochord development. Both structures derive from an early population of axial mesodermal progenitor cells. Cells that eventually become the notochord require *foxa2*, *foxa3*, *noto*, and *tbxta*, whereas *tbxta* is not required for the specification differentiation of axial mesodermal cells into the PC. (notochord is depicted in green and PC is depicted in purple at 3 dpf when PC is maturing).

